# Monocyte-derived macrophages promote intraepithelial infiltration of effector memory CD8^+^ T cells in tumors regressing after STING agonist treatment

**DOI:** 10.64898/2026.05.28.728475

**Authors:** Anaïs Vermare, Annavera Ventura, Adrien Rouault, Luca Simula, Malvina Seradj, Leïla Lhuillier, Eleonore Weber-Delacroix, Kevin Mulder, Lene Vimeux, David Espié, Karine Bailly, Brigitte Izac, Benjamin Saintpierre, Wail Zeitouni, Ariane Jolly, Pierre Delagrange, Marion V Guérin, Emmanuel Donnadieu, Frederic Pendino, Charles-Antoine Dutertre, Alexandre Boissonnas, Armelle Prévost-Blondel, Elisa Peranzoni, Nadège Bercovici

## Abstract

Despite the clinical success of cancer immunotherapies, the cellular interactions driving tumor regression remain incompletely understood. Here, we investigated the dynamic remodeling of the tumor immune microenvironment during regression of transplanted PyMT mammary tumors following STING agonist treatment. Using scRNA-seq of sorted CD8^+^ T cells and myeloid cells, combined with imaging approaches, we identified major changes in both lymphoid and myeloid compartments during tumor regression. Regressing tumors showed a transient accumulation of Ly6C^hi^ monocyte populations associated with a decline in macrophage subsets, while effector and memory CD8^+^ T-cell populations increased at the expense of exhausted T cells. Interaction analyses predicted enhanced chemotactic and adhesion interactions between CXCL9^+^ Ly6C^hi^ monocytes and effector CD8^+^ T cells. Consistently, dynamic imaging revealed increased CD8^+^ T-cell motility and infiltration into tumor cores following treatment. In particular, CXCR6^+^ effector CD8^+^ T cells transiently accumulated within tumor islets during regression before relocalizing to stromal regions. Together, these findings reveal a coordinated spatiotemporal remodeling of myeloid and CD8^+^ T-cell populations during immunotherapy-induced tumor regression and highlight cooperative interactions that may promote durable anti-tumor immunity.

## INTRODUCTION

Tumors are complex ecosystems, made up of various cell types, including lymphocytes and myeloid cells, in addition to cancer cells themselves. All are intertwined in complex networks of interactions. During tumor progression, the cancer cells are known to polarize myeloid cells, which are particularly plastic, into pro-tumoral actors through systemic secretions of various cytokines and growth factors, as reviewed in (Mantovani et al. 2002). Consequently, macrophages at the tumor site are, for the great majority, M2-like tumor-supporting cells, harboring immunosuppressive properties against anti-tumoral effector CD8^+^ T cells.

Some immunotherapies, such as immune checkpoint blockade, target specific receptor/ligands such as PD1, PDL1, CTLA-4 on cells to lift off the inhibitory interactions of effector CD8^+^ T cells with the cancer cells and immunosuppressive cells. Other therapies like radiotherapy or stimulator of interferon genes (STING) agonists, rather affect the system as a whole, through the induction of pro-inflammatory pathways, for instance type I interferons (IFN). Strikingly, in both cases, the immunotherapy treatment remodels more than just the initially targeted CD8^+^ T cell compartment. For example, some cancer types, including breast cancer, treated with immune checkpoints blockade, a modification of the myeloid compartment has been reported, in addition to the directly targeted T cell populations (Bassez et al. 2021; Gubin et al. 2018).

This implies there will be changes of interactions between myeloid cells and CD8^+^ T cells in tumors post-immunotherapy, which some studies already suggest. Indeed, myeloid cells and CD8^+^ T cells interactions in tumors were classically thought to be deleterious, as synthesized in (Mantovani et al. 2017). However, various reports have shown that macrophages and monocytes monocyte-derived macrophages can actually cooperate with anti-tumoral CD8^+^ T cells when appropriately polarized by therapeutic interventions. Our team and others have notably demonstrated that the presence of both CD8^+^ T cells and macrophages is necessary for an optimal tumor regression post-immunotherapy (Thoreau et al. 2015; Weiss et al. 2017).

The use of a STING agonist requires downstream secretion of type I IFNs by activated MHCII^+^ TAMs to induce tumor regression (Guerin et al. 2019).Type I IFNs drive pro-inflammatory polarization of monocytes and macrophages in various models (Kwart et al. 2022; Lam et al. 2021) and immunotherapy efficacy was furthermore associated to these alternatively-polarized myeloid cell presence, highlighting once more the need to re-evaluate myeloid cells functions after treatment.

Yet, the literature on the positive impacts of activated myeloid cells towards T cells is quite recent and the exact functions and interactions between those actors remain to be understood, in order to be exploited to increase immunotherapy efficacy.

In this study, we therefore investigated the mechanisms by which macrophages and monocytes communicate with effector CD8^+^ T cells at the tumor site during tumor regression. We used the MMTV-PyMT breast cancer model in which we previously showed macrophages and CD8^+^ T cell cooperate after a single i.p. injection of a STING agonist. We performed single-cell RNAseq on both cell types and revealed an increase in the monocyte-to-macrophage ratio in tumors after treatment, as well as an enrichment in memory-like populations of CD8^+^ T cells. *In silico* predictions highlighted chemotaxis activity between inflammatory monocytes and effector/effector memory CD8^+^ T cells, and indeed, overall intra-epithelial infiltration and migration of CD8^+^ T cells was increased after STING agonist treatment. Furthermore, the presence of such inflammatory monocytes is also predictive of good response to immunotherapy in breast cancer patients.

## RESULTS

### Effectors and stem-cell memory CD8^+^ T cells are enriched in regressing tumors after STING agonist treatment

The treatment with DMXAA induced the regression of PyMT mammary tumors **(Fig. 1A)**, leading to immune protection with a prolonged survival in the majority of the mice **(Fig. S1)**. To characterize the tumor-infiltrating CD8^+^ T lymphocytes (TILs) in regressing tumors, tumors were collected 4 days after DMXAA treatment, when tumors started to regress, and CD8^+^ TILs were isolated **(Fig. 1B and 1C)**. The treatment induced the progressive accumulation of CD8^+^ TILs compared to progressing tumors, reaching almost 40% of CD45^+^ cells within 8 days **(Fig 1D)**.

**Figure 1.**
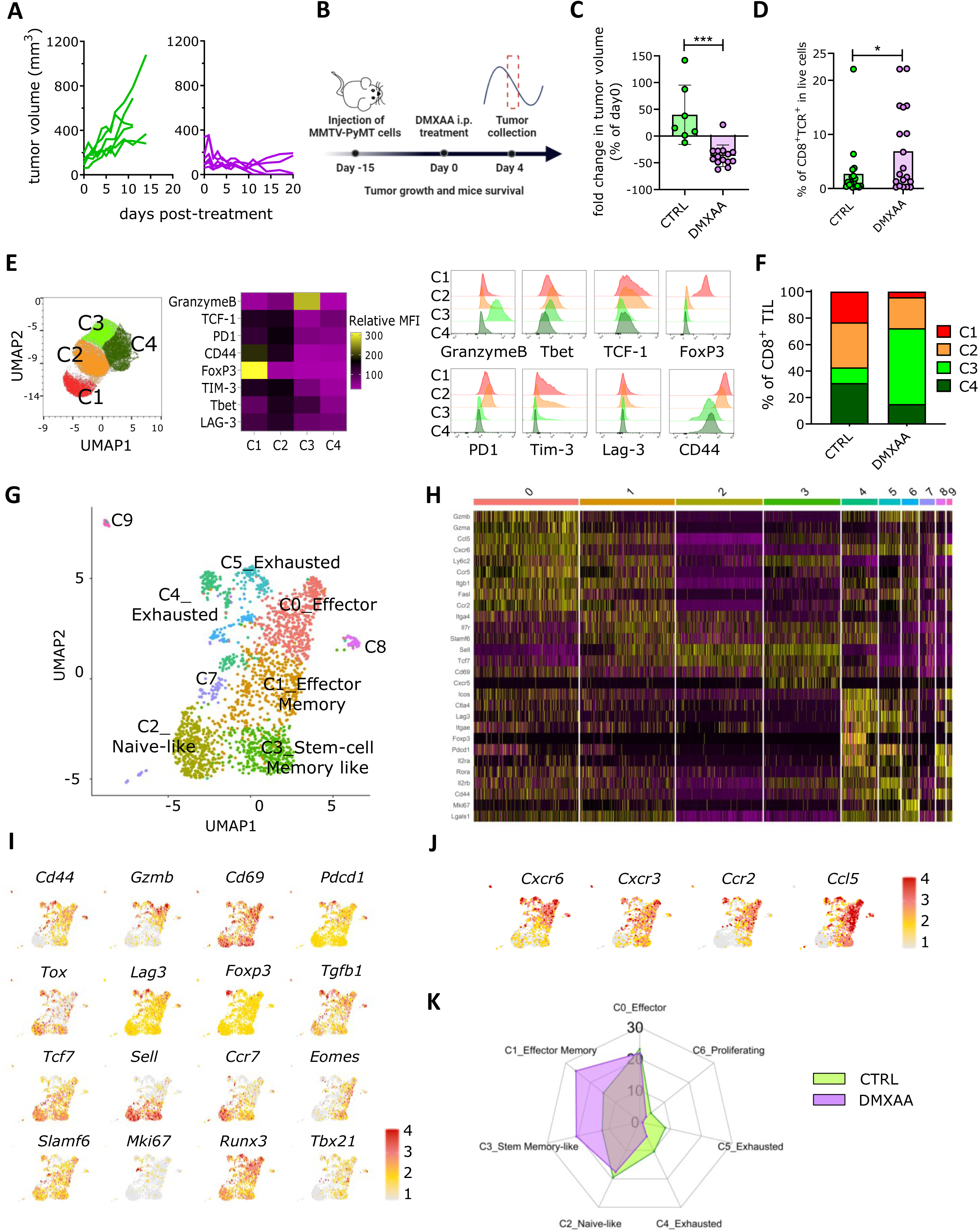
Enrichment of effector and memory subsets of CD8^+^ TILs in breast cancer tumors post-STING agonist treatment. **(A)** FvB mice transplanted with PyMT tumors were treated at day 0 with one i.p injection of DMXAA (purple graph) or PBS 50% DMSO as a control (green graph), two weeks post-transplantation as explained by the cartoon in **(B)**. **(C)** The fold change of tumor volumes at day 4, relative to day 0 are represented for control and treated groups. DMXAA, n=14 and controls n=7, from 4 independent experiments. **(D)** The percentage of CD8^+^ T cells among CD45^+^ cells in control (green), day 4 and day 8 (purple) tumors was determined by flow cytometry. DMXAA, n= 19 and controls n=18, from 4 independent experiments. Kruskal-Wallis multiple comparison non parametric test. * p < 0.05; *** p < 0.001. **(E)** CD8^+^ TIL diversity was assessed by spectral flow cytometry through the expression of surface and intracellular markers. The UMAP represents the subset clusters obtained by analyzing all the CD8^+^ TIL, independently of their tumor of origin. The heatmap summarizes the relative mean fluorescence intensity (MFI) of each analyzed marker between the four subsets C1, C2, C3 and C4, as illustrated by the histograms on the right. **(F)** Proportions of each of the four clusters in CD8^+^ TILs according to control or day 4 treated tumors are represented in bar histograms. DMXAA, n=2 and controls n=2, from one experiment. **(G)** UMAP plot of single-cell RNAseq (scRNAseq) analysis of sorted CD8^+^ TIL from merged control and day 4 treated tumors, with clusters labels proposed upon **(H)** clusters selected top genes heatmap and **(I)** signature genes expression UMAPs examination. (J) Radar plot depicting cluster percentages in CD8^+^ TILs from control or day 4 treated tumors. DMXAA, n=26 and controls n=6, merged by group for one experiment.

Single-cell analysis of control and treated mice by spectral cytometry segregated four subsets of CD8^+^ TILs **(Fig. 1E)**. Two clusters, C1 and C2, expressed high levels of the inhibitory receptors PD-1, TIM-3 and LAG-3, but C1 expressed the transcription factor FOXP3 while C2 expressed the Th1 transcription factor TBET. By contrast, cluster C3 expressed high levels of granzyme B and a fourth cluster C4 expressed none of these effector molecules. When comparing the profiles of CD8 TILs in treated and control mice, we observed that the cluster C3 of GrzB^+^ CD8^+^ TIL were enriched in regressing tumors, while the FOXP3^+^ C1 cluster was drastically reduced **(Fig. 1F)**. These data indicate that the STING agonist treatment rapidly induces a reduction of exhausted CD8^+^ TILs and promotes the infiltration of GZMB+ effector CD8 TIL.

To increase our resolution of CD8^+^ TIL subsets infiltrating regressing PyMT tumors, we performed single-cell RNA sequencing (scRNAseq) on sorted CD8^+^ TILs. Uniform manifold approximation and projection revealed ten distinct subsets of CD8^+^ TILs **(Fig. 1G and 1H)**: effector CD8^+^ TILs with Gzmb^+^ Effector T cells (cluster C_0) and Slamf6^+^effector-memory T cells (cluster C_1), less differentiated CD8^+^ TILs with Lef1+ Tcf7+ T cells (cluster C_2), Tcf7^+^ progenitor–exhausted T cells (cluster C_3) and populations of highly differentiated, exhausted CD8^+^ T cells expressing Foxp3 (cluster C_4) or Pdcd1 (cluster C_5). We also identified Mki67+ proliferating T cells (cluster C_6) and three minor clusters of CD8^+^ T cells expressing Treg and innate-like markers such as Cd74 (Cluster C_7), Xcl1(Cluster C_8) and Ramp1^+^ (Cluster C_9) **(Fig. S2)**. The population of effector/effector-memory and exhausted CD8^+^ TILs all expressed the markers of residency Itgae or Cxcr6 **(Fig. 1I)**. Interestingly, the exhausted CD8^+^ TIL clusters C_4 and C_5 was greatly reduced in DMXAA-treated mice compared to control mice. In sharp contrast, regressing tumors were enriched in effector-memory T cells (cluster C_1) and progenitor–exhausted T cells (cluster C_3) (Fig. 1J). These two populations together with the C_0 effector CD8^+^ TIL cluster represented the majority of CD8^+^ TILs in regressing tumors after STING agonist treatment. These data confirmed our spectral cytometry findings and suggest that the STING agonist treatment can re-activate the CD8^+^ TIL compartment.

### Interferon reprograms CD8^+^ TIL and foster the intraepithelial infiltration of CXCR6^+^ CD8^+^ TIL into tumor islets

To get further insights into the CD8^+^ TIL subsets activated by the STING agonist therapy, we performed a gene ontology analysis in CD8+ TIL clusters in treated versus control mice **(Fig. 2A)**. We found that production and response to type I interferon were mainly restricted to Lef1^+^ Tcf7^+^ cluster C_2 and to Tcf7^+^ C_3 and Slamf6^+^ effector-memory C_1 clusters, respectively, suggesting that these were the primary target of the STING agonist. Upregulation of pathways related to T cell activation, differentiation and IFNγ production were found in several CD8^+^ TIL clusters.

**Figure 2.**
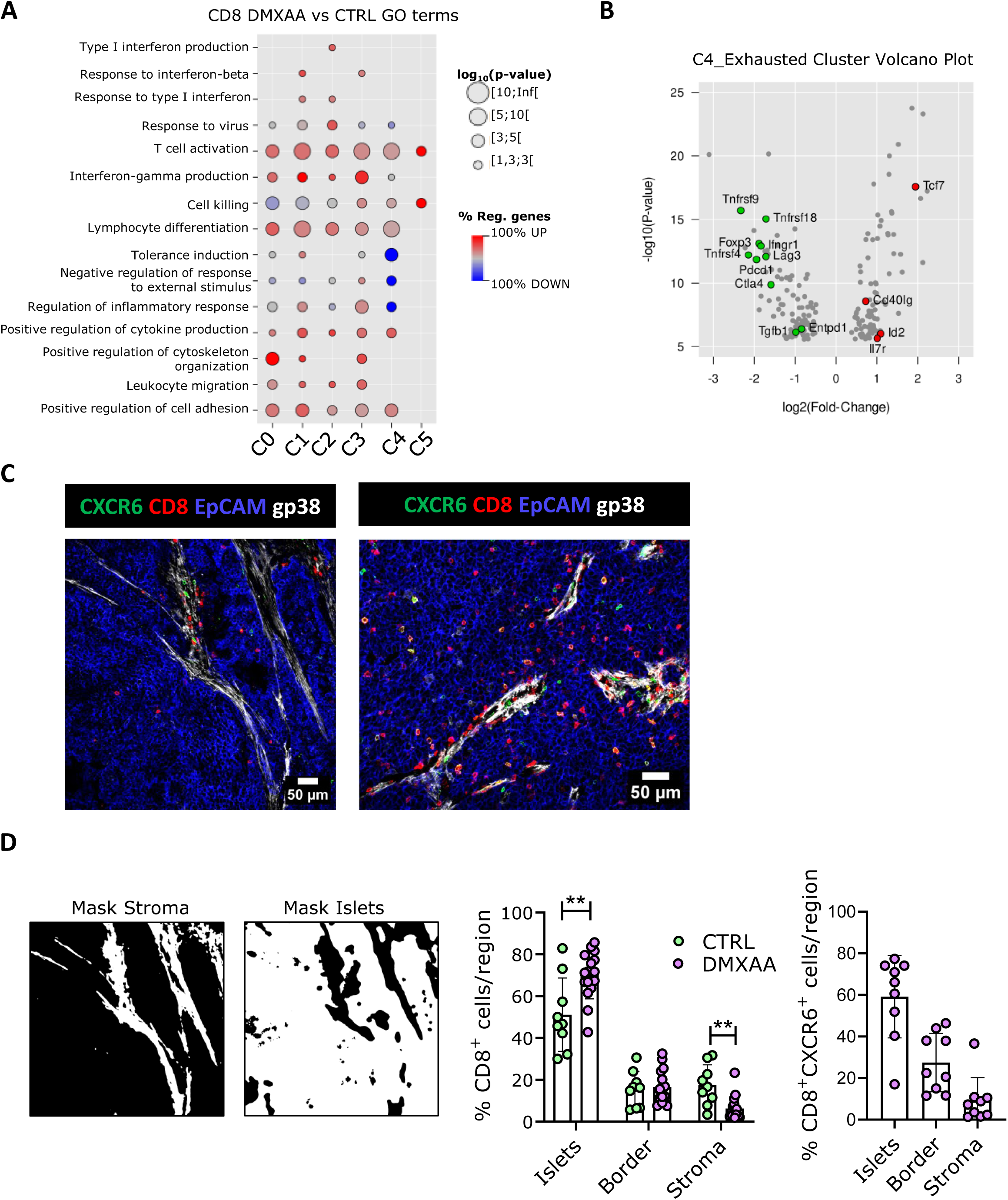
Enhanced activation and tumor islet infiltration by CD8^+^ TILs after treatment with DMXAA. **(A)** Upregulated (red) and downregulated (blue) pathways in the scRNAseq CD8^+^ TIL clusters after mice DMXAA-treatment were determined using Gene Ontology (GO) analysis and displayed on a bubble plot, with p-value significance represented by bubble size. **(B)** The volcano plot highlights the up (red) and down (green) regulated genes within the cluster 4 of CD8^+^ TILs from treated tumors compared to controls. **(C)** Representative images to illustrate the positioning of CXCR6^+^ or CXCR6^neg^ CD8^+^ T cells in tumor compartments (EpCAM^+^ tumor islets, blue and gp38^+^ stroma, grey) in control tumors and day 4 or 6 treated tumors. Maximum intensity Z projection visualization with 20x magnification, scale bar of 50 µm. **(D)** The percentage of CD8^+^ TIL or CXCR6^+^ CD8^+^ positioning in tumor compartments (islets, stroma and border (overlap of islet and border stainings)) was quantified with the help of an algorithm specifically-developed, using masks to delineate between tumor regions, as shown on the left. Quantification results are expressed as mean ± SEM. DMXAA (day 4 and 6) n=5 mice and controls n=4, with at least two images per tumor, from 4 or 5 independent experiments. ** p < 0.01.

Interestingly, the Foxp3^+^ T cell cluster C_4 showed a strong down regulation of tolerance induction pathways. We further looked at the genes differentially expressed in this Foxp3^+^ C_4 cluster between treated and control mice **(Fig. 2B)**. We found that indeed, the STING agonist treatment led to a strong downregulation of Foxp3, Tgfb1 and inhibitory receptors Pdcd1, Ctla4 and Lag3, all involved in immunosuppressive T-cell functions. The expression of costimulatory molecules of the TNFR superfamily (Tnfrsf4, Tnfrsf9, Tnfrsf18), were also down-regulated. In contrast, this subset of CD8^+^ T cells upregulated Tcf7 and Id2, involved in the transition to exhausted progenitors and terminally exhausted TILs. These findings indicate that type I IFN reprograms CD8^+^ T cell gene expression profiles with the emergence Tcf7^+^ progenitors and effector-memory CD8^+^ TILs.

We next examined the impact of this reprogramming on the capacity of CD8^+^ T cells to infiltrate the tumor compartment. The gene ontology analysis indeed revealed the upregulation of various genes involved in leukocyte migration and cell adhesion **(Fig. 2A)**. Although intraepithelial infiltration of CD8^+^ T cells can rarely be assessed in mouse models because most tumor cell lines have undergone an epithelial-to-mesenchymal transition losing their distinctive epithelial markers, we took advantage of a PyMT mouse model that nicely recapitulates the architecture found in human carcinomas (Guerin, elife 2019) to track the infiltration of CD8^+^ TILs during tumor regression. We performed multiplex immunofluorescence microscopy on tumor slices from control and STING-agonist treated mice **(Fig. 2C)** and we used the CXCR6 marker to localize the clusters of effector/effector-memory CD8^+^ TILs expanded after treatment. Whereas CD8^+^ TILs are mostly found in the stromal area of the tumor mass in control mice, these cells were enriched in tumor islets after STING agonist treatment **(Fig 2D)**. Quantification of CXCR6^+^ CD8^+^ TILs further showed that the majority of resident effector-memory CD8^+^ TIL indeed reached tumor islets. Together, these data show that the tumor regression induced by STING agonist treatment is associated with an intraepithelial infiltration of re-activated CD8^+^ TIL subsets, including resident memory CD8^+^ TILs.

### CD8^+^ TIL infiltration of regressing tumors is accompanied by remodeling of the macrophage compartment

As we previously showed that tumor-associated macrophages (TAMs) constitute a major obstacle to intraepithelial T cell infiltration, we wondered if STING agonist treatment had modified the myeloid compartment in such a way that T cells were able to better infiltrate the tumor compartment. To address this question, we phenotypically characterized tumor-infiltrating mono-macrophages in control and regressing tumors by spectral cytometry **(Fig. 3A and Fig. S3)**. We distinguished three populations of TAMs (F4/80^+^ CD64^+^ immature TAMs, CX_3_CR1^+^ PDL1^neg^ TAMs and CX_3_CR1^+^ PDL1^+^ TAMs) as well as CD64^+^ monocytes (TAMo), classical monocytes (cMo) and non-classical monocytes (ncMo). Strikingly, early after STING agonist treatment, we observed a drastic reduction of TAMs paralleled by an accumulation of monocytes in regressing tumors **(Fig. 3A and 3B)**. The reduction of TAMs in treated mice reverted the monocyte-to-TAM ratio, reaching an average of 6 monocytes/TAMs compared to a ratio of 0.35 in control mice. This phenomenon was however transient as the TAM population had almost completely recovered when tumors were analyzed at day 8 after treatment **(Fig. 1A)**. We hypothesized that the inversion of the monocyte-to-TAM ratio was correlated with tumor regression. As we previously reported that MMTV-PyMT mice that spontaneously develop mammary tumors were resistant to STING agonist treatment (Guerin et al. 2019) we analyzed the monocyte-to-TAM ratio in MMTV-PyMT mice 4 days after treatment with the STING agonist **(Fig. 3C)**. Despite a significant increase in the monocyte-to-TAM ratio, TAMs still outnumbered monocytes in these resistant tumors.

**Figure 3.**
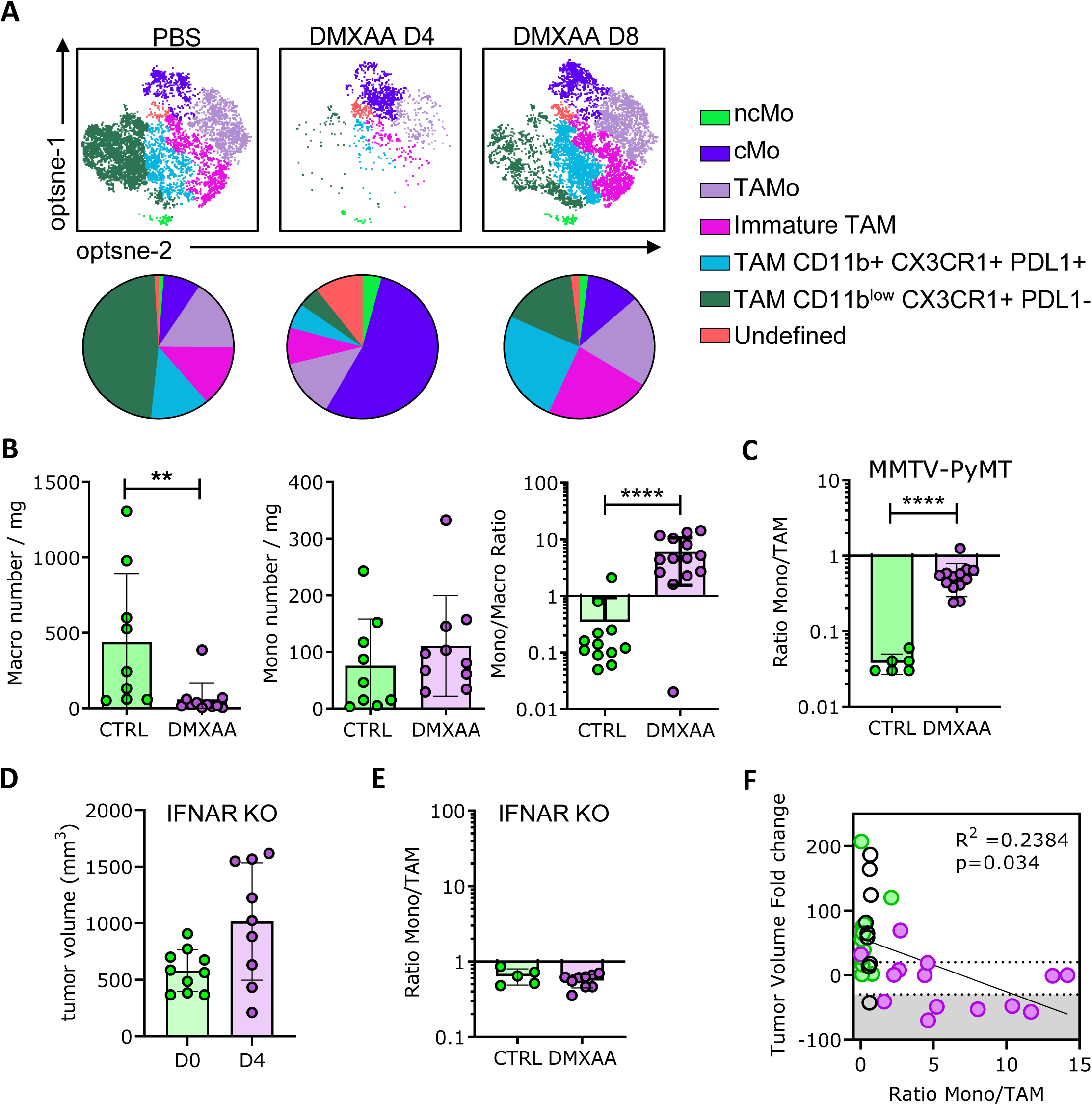
Transient type I IFN signaling-dependent replacement of tumor-associated macrophages by inflammatory monocytes in mice tumors treated by DMXAA. **(A)** Monocyte and macrophages diversity was assessed by spectral flow cytometry and divided into clusters, named after their markers expression signature (ncMo: Ly6C^neg^, CD43^+^, cMo: Ly6C^+^ CCR2^int^, TAMo: Ly6C^+^ CD64^+^ MHCII^+^, Immature TAM: CD64^int^ F4/80^int^ PDL1^+^, TAM: CX3CR1^+^ F4/80^+^ CD64^+^. (The undefined population did not fit in any of the previously detailed subpopulations). The tSNE represents the clusters present in control, day 4 and day 8 treated mice tumors and frequencies of each subpopulation for each condition are represented in color-coded pie charts. **(B)** Absolute numbers of monocytes (Ly6G^neg^ Ly6C^+^), macrophages (Ly6G^neg^ Ly6C^neg^ F4/80^+^) and monocyte-to-macrophage ratio were determined by flow cytometry in control (green) or day 4 treated tumors (purple). DMXAA, n=8 and controls n=6, from 3 independent experiments. Mann-Whitney statistical test. ** p < 0.01, **** p < 0.0001. **(C)** day 4 DMXAA, ratio % in MMTV-PyMT tumors. **(D)** IFNAR^-/-^ PyMT tumors were transplanted in C56BL/6 mice and treated with DMXAA. Tumor volume is represented at day 0 and day 4 post-treatment, as well as **(E)** monocyte-to-macrophage ratio obtained through flow cytometry experiments at day 4. Controls n=5, DMXAA n=9 mice, from one experiment. **(F)** Monocyte-to-macrophage ratio represented against tumor volume foldchange (day 4 relative to day 0). Data is a merge from control (green) and treated (purple) mice, as well as from IFNAR^-/-^ tumor experiments. Correlation analysis, r^2^=0.2384 with * p < 0.05.

We then wondered if the drastic decrease in TAMs in the sensitive transplanted PyMT mice was linked to IFN type 1. Indeed, we previously showed that tumor regression induced by STING agonist was dependent on the expression of type 1 IFN receptor (IFNAR) by host cells and that the resistance of MMTV-PyMT mice was partly due to the inhibition of type 1 IFN production by TGFβ (Weiss et al. 2017). To address this question, we treated IFNAR KO transplanted PyMT mice with the STING agonist and looked at both tumor regression and monocyte-to-TAM ratio. Tumor regression was completely blocked in these mice, with no change in the monocyte-to-TAM ratio **(Fig. 3D and 3E)**. These findings suggest that TAM depletion following STING agonist treatment depends on type 1 IFN. Furthermore, we found that the monocyte-to-TAM ratio was directly correlated with tumor regression **(Fig. 3F)**.

Taken together, these results indicate that STING agonist treatment promotes a drastic remodeling of the macrophage compartment, with a window of acute inflammation favoring the accumulation of newly-recruited monocytes over TAMs in regressing tumors.

### Monocytes with anti-tumoral properties infiltrate regressing tumors

We next characterized the populations of mono-macrophages reprogrammed by the STING agonist treatment by scRNAseq **(Fig. 4A and 4B)**. We identified six clusters of macrophages, including Cx3cr1^+^ TAMs (C_0), Ftl1^+^ TAMs (C_1), Mki67^+^ TAMs (C_2 and C_6), Ifit^+^ TAMs (C_3), Mmp9^+^ TAMs (C_7), two clusters of monocytes, Cxcl9^+^ monocytes (C_4) and non-classical Nr4a1^+^ monocytes (C_8)), and one cluster of Ccr7^+^ dendritic cells (DCs) (C_5).

**Figure 4.**
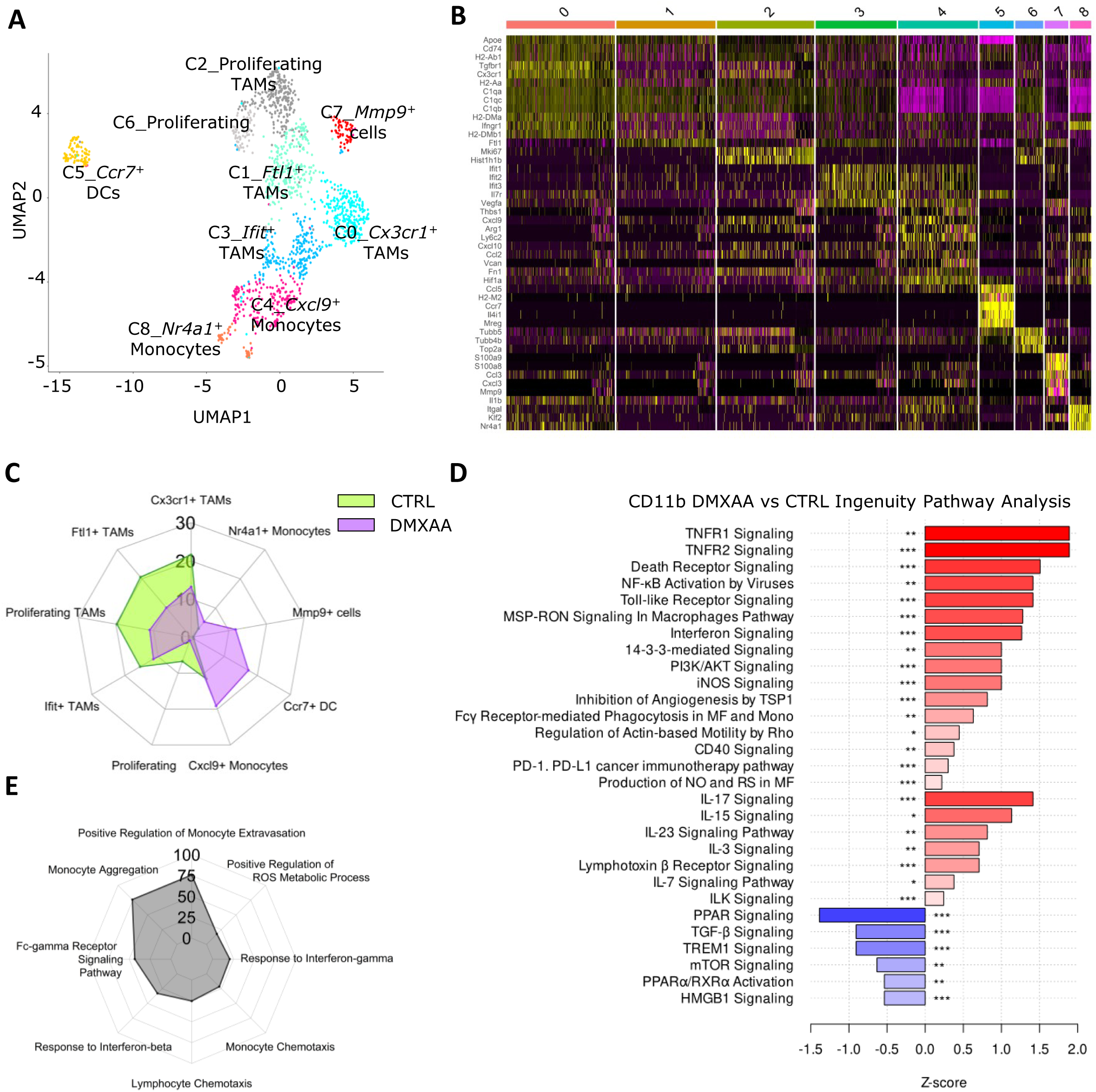
Upregulation of CD8^+^ T cell response supporting functions in inflammatory monocytes freshly recruited at the tumor site. **(A)** UMAP plot of scRNAseq analysis of sorted monocyte and macrophages populations (CD11b^+^ NK1.1^neg^ Ly6G^neg^) from merged control and day 4 treated tumors, with clusters labels proposed upon **(B)** clusters top selected genes heatmap examination. **(C)** Radar plot showing cluster percentages among monocyte/macrophage population in control or day 4 treated tumors. DMXAA, n=26 and controls n=6, merged by group for one experiment. **(D)** Upregulated (red) and downregulated (blue) pathways in the scRNAseq whole monocytes/macrophages population after mice DMXAA-treatment were determined using Ingenuity software and selected pathways of interest represented according to their Z-score and p-value (* p < 0.05, ** p < 0.01, *** p < 0.001). **(E)** Selected predicted functions for cluster 4 of the monocyte/macrophage population obtained through GO analysis, represented in terms of fold enrichment.

Consistently with the spectral cytometry data, the TAM population collapsed in regressing tumors **(Fig. 4C)**, with five TAM clusters reduced. Conversely, an enrichment in monocytes (Cxcl9^+^ C_4 and Nr4a1^+^ C_8), Ccr7^+^ DCs (C_5) and Mmp9^+^ TAMs (C_7) was observed. These data indicate that by depleting the TAM niche, STING agonist treatment creates a window of acute inflammation resulting in the accumulation of new populations of myeloid cells at the tumor site.

To get an idea of the reorientation of the myeloid profile in tumors after treatment with the STING agonist, we next used ingenuity pathway analysis to compare the predicted functions of myeloid cells in regressing tumors compared to control tumors. Several pathways related to macrophage activation and phagocytosis were upregulated after treatment, whereas TGFβ signaling and metabolic regulator of macrophage activity (mTOR) were downregulated **(Fig. 4D)**. These data show that STING agonist treatment reshapes the myeloid compartment towards a less immunosuppressive and more anti-tumoral myeloid cell profile.

To understand the biological functions of different myeloid cell clusters and their potential influence on CD8^+^ TIL infiltration, we performed Gene Ontology analysis on clusters enriched following STING agonist treatment. We concentrated on the monocyte cluster C_4, which expresses the T cell chemoattractant CXCL9, since the density of CXCL9^+^ cells in tumors is crucial for T cell infiltration (Dangaj et al. 2019). Notably, we discovered an enrichment of genes involved in the response to type I interferon, reactive oxygen species (ROS) and nitric oxide (NO) production, cell migration, extravasation and the regulation of lymphocyte chemotaxis **(Fig. 4E)**. These results led us to further investigate if these cells could play a role in the progressive infiltration of CD8^+^ TILs within epithelial tumor cell islets during regression.

### STING agonist treatment fosters CD8^+^ TIL-myeloid cell cooperation for cell migration

First, we analyzed the motility of CD8^+^ TILs in the 3D tumor microenvironment of fresh tumor slices by live imaging **(Fig. 5A)**. CD8^+^ TILs infiltrating tumors at the early stages of regression (4 to 6 days) showed an increased average speed and distance traveled compared to control tumors. We could observe these cells moving across the intraepithelial EpCAM^+^ area (Supplementary Movie 1), indicating that they were activated and highly motile. Surprisingly, they rarely stopped except in the proximity of myeloid cells found in these areas. Here, they made long-lasting interactions with myeloid cells during the recording period **(Fig. 5B)**. In control tumors, CD8^+^ TILs were rare but some interactions with myeloid cells in stromal regions were observed, in line with our previous report (Peranzoni et al. 2018). These observations indicate that despite contacting myeloid cells, CD8^+^ TILs migrated more actively in regressing tumors compared to progressing ones.

**Figure 5.**
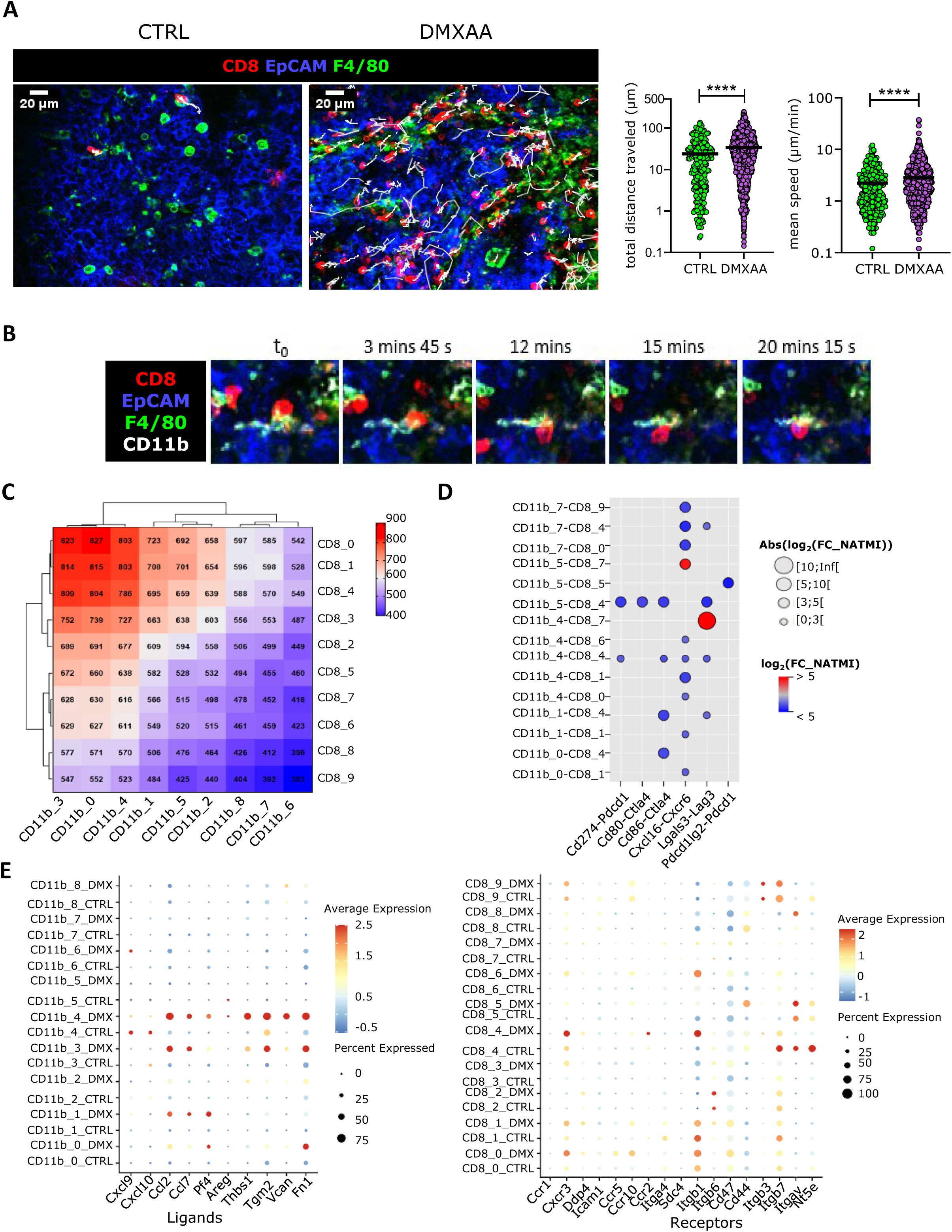
Migration-enhancing interactions predicted between effector CD8^+^ T cells and monocytes in tumors post-DMXAA treatment. **(A)** Live imaging movies following CD8+ cells migration in slices of tumors, with cell mean speed (µm/min) and distance traveled (µm) quantifications, from control or treated mice (day 4-6 post-DMXAA). DMXAA, n=6 tumors and controls n=4 tumors, with a minimum of one slice per tumor, from 4 independent experiments. Statistical test and * p value to add when D4 and D6 data are merged. **(B)** Series of snapshots showing a CD8^+^ T cell making long-lasting interactions with a F4/80^+^ macrophages at the tumor site. **(C)** Heatmap visualization of the number of predicted interactions on the basis of the scRNAseq data analyzed in three interaction prediction algorithms (NATMI, CellPhoneDB and NichNet), between cells from each CD11b^+^ clusters and each CD8^+^ T cell clusters. **(D)** Up (red) and down (blue) regulated ligand-receptor interaction pairs between specific CD11b^+^ and CD8^+^ clusters in treated tumors compared to controls using the NATMI algorithm. Bubble size and color represent the interaction foldchange. **(E)** Ligands (on CD11b^+^ cells) and receptors (on CD8^+^ T cells) of interest were selected and their relative expression was represented on bubble plots. Each cluster was re-divided between control and treated origins to show the average expression of the ligand/receptor in the given cluster and condition (bubble color) as well as the percentage of cells actually expressing the gene (bubble size).

To decipher the nature of the interactions taking place between myeloid cells and CD8^+^ TILs in this context, we explored the interactions predicted to occur between these cell subsets based on scRNAseq data. Most of the interactions involved the effector/effector-memory (C_0 and C_1) and exhausted C_4 CD8^+^ TIL clusters on one hand and the Cx3cr1^+^ TAMs (C_0), Ifit^+^ TAMs (C_3), and the Cxcl9^+^ C_4 monocytes clusters on the other **(Fig. 5C)**.

When we compared how these interactions were differently regulated between regressing and progressing tumors **(Fig. 5D)**, we observed that several interactions involving inhibitory receptors Pdcd1, Ctla4 and Lag3 on exhausted CD8^+^ TILs and their ligands on Ccr7^+^ DCs, TAMs and monocytes were decreased in regressing tumors. Only the Lag3/Lgals3 axis was specifically increased between the C_7 CD8^+^ TIL cluster and monocytes. Interestingly, Pdcd1 (PD-1) on CD8**^+^** TIL was almost exclusively found to interact with Cd274 (PD-L1) expressed on Ccr7^+^ DCs, whereas Pdcd1lg2 (PD-L2) on Mmp9**^+^** TAMs engaged PD-1 on exhausted CD8^+^ TILs (C_5). These data show that the reprogramming induced by the STING agonist treatment drastically decreases the engagement of immune checkpoints inhibitors by myeloid cells.

Moreover, we found that interactions through the Cxcl16/Cxcr6 axis engaged by TAMs and Cxcl9^+^ monocytes were decreased in regressing tumors. These results suggest that additional chemokines might favor the migration of Cxcr6^+^ CD8^+^ TILs into tumor islets during regression **(Fig. 2C and 2D)**.

We then looked at the interactions that were upregulated during tumor regression between CD8^+^ TILs and myeloid clusters, in particular with the CXCL9^+^ monocytes cluster C_4 that seems to mediate lymphocyte chemotaxis. We found that the top predicted interactions with this cluster were chemokines and adhesion molecules. Unexpectedly, CCL2, whose expression was strongly upregulated after treatment, was predicted to engage CCR2 on effector/effector memory CD8^+^ TILs **(Fig. 5E, left panel)**. The expression of CCR2 was also upregulated on CD8^+^ TILs **(Fig. 5E, right panel)**. In contrast, we did not find evidence for interactions occurring through the Cxcl9/10-Cxcr3 axis. Indeed, the treatment increased the expression of Cxcr3 on CD8^+^ TILs, but was accompanied by a decreased expression of Cxcl9/10 in Cxcl9^+^ monocytes. These data indicate that despite high expression of Cxcl9 and Cxcl10 at baseline, these monocytes upregulate additional chemo-attractants in regressing tumors.

We sought to validate experimentally the involvement of CCL2 produced by Cxcl9^+^ monocytes in CD8^+^ TIL infiltration by targeting the CCR2/CCL2 axis. First, we confirmed that CCL2 was produced and secreted in regressing tumors **(Fig. 6A)** and that CD8^+^ TILs upregulated CCR2 expression at their cell surface **(Fig. 6B)**. Analysis of CD8^+^ TIL behavior by live imaging showed that the motility of CD8 TIL from STING agonist treated mice was decreased to control levels in some tumor samples after in vitro CCR2 blockade **(Fig. S4)**. These data suggest that the CCR2/CCL2 axis may play a role early in tumor regression to support the recruitment and infiltration of CD8^+^ TILs in tumors. Together, these findings reveal a dynamic network of interactions between effector/effector memory CD8^+^ TILs and Cxcl9^+^ monocytes at an early stage of tumor regression.

**Figure 6.**
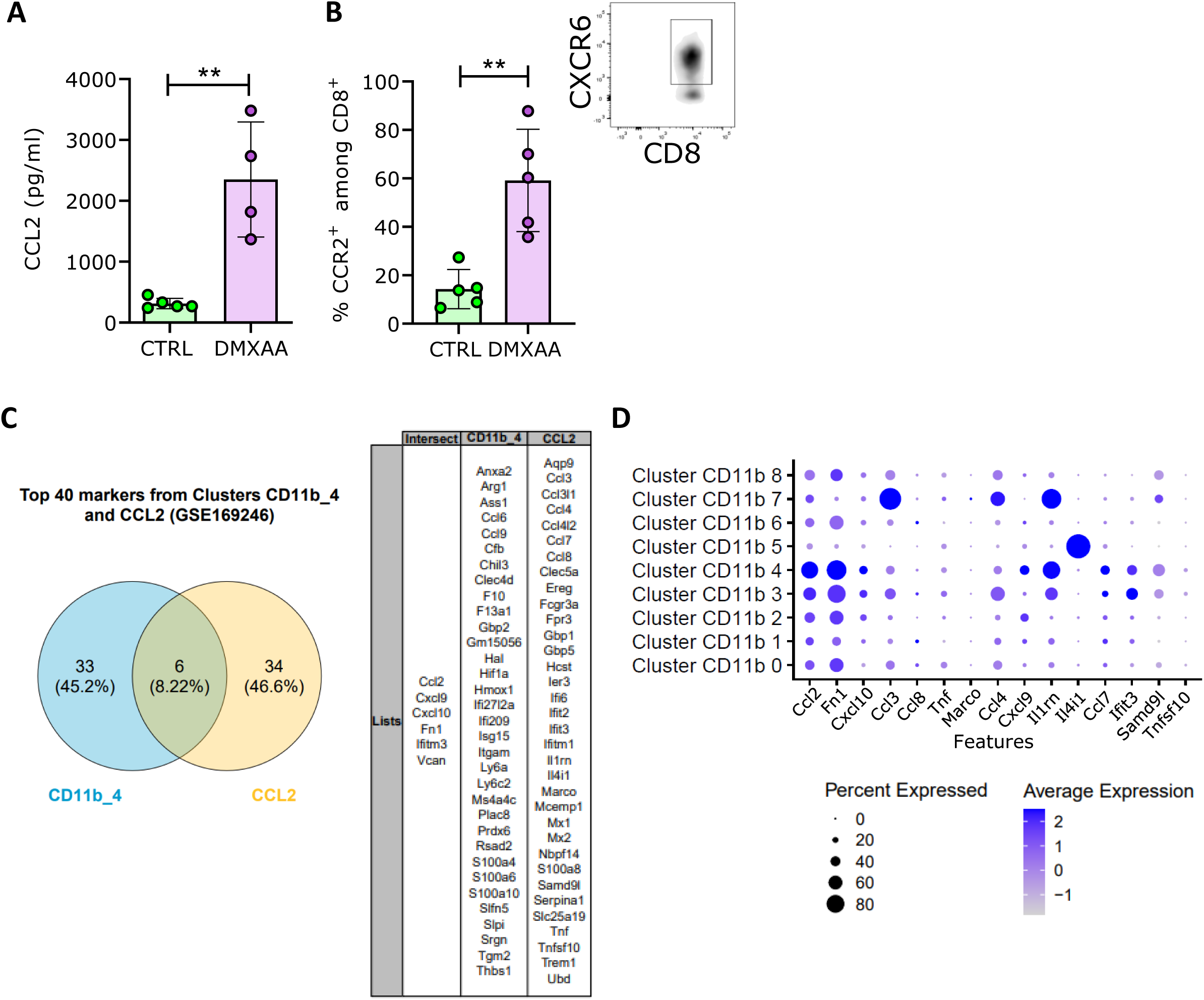
Predicted CCL2-CCR2 axis is increased after treatment in our model of breast cancer and is found to be of good prognosis in human data. **(A)** CCL2 production by control and day 4 treated tumors was assessed by dosing the supernatants of tumor slices after a 48 hours incubation. DMXAA, n= 4 and controls n= 5, from 2 independent experiments. T-test statistical test. ** p < 0.01. **(B)** The percentage of CD8+ T cells expressing CCR2 was determined by flow cytometry, in control and day 4 treated mice tumors. T-test statistical test was performed, ** p < 0.01. **(C)** Comparison of top 40 markers between cluster 4 of CD11b^+^ from our work and Zhang’s study t-macro-CCL2 subset is represented in a Venn diagram for the number of intersect and individual genes. **(D)** Expression of the top 15 signature genes of t-macro-CCL2 (Zhang’s study) in CD11b^+^ clusters from our work. The bubble plot allows for the representation of each gene average expression in cluster (bubble color) and the percentage of population expressing it (bubble size).

### CXCL9^+^ CCL2^+^ monocyte/macrophage population infiltrates human tumors and is a predictive marker of response to immunotherapy

To further investigate the relevance of our findings in human cancer, we investigated if TAM subsets similar to those we found enriched in regressing tumors had been described inhuman tumors. We used the MoMAC-Verse single-cell Atlas of human monocytes and macrophages from human tissues (Barras et al. 2024; Mulder et al. 2021) to identify the human ortholog of our myeloid clusters **(Fig. S5A and 5B)**. The mouse Cxcl9^+^ monocytes were found to project onto the IL4I^+^ macrophage population, with Cxcl9 and Cxcl10 in the top 40 differentially expressed genes. This CXCL9^+^ macrophage population was described to interact closely with T cells in the periphery of human tumor tissues (Mulder et al. 2021)during immunosurveillance. However, the impact of immunotherapy on this network of interactions remains to be discovered. The data we obtained in our PyMT mouse model suggest that the acute inflammation induced by immunotherapy may stimulate chemotaxis through CCL2 expression by this Cxcl9^+^ macrophage population, promoting infiltration of effector/effector memory CD8^+^ TILs. In human breast cancer, we found that Zhang and colleagues described the existence of macrophages expressing high levels of CCL2 and CXCL9/10 that were predictive of response to the combination of anti-PD-L1 and chemotherapy (Zhang https://doi.org/10.1016/j.ccell.2021.09.010, 2021) (Zhang et al. 2021). By mapping our scRNAseq on this data set, we found many similarities between our Cxcl9^+^ monocytes and this human CCL2-macrophage population **(Fig. 6C)**, as they both express high levels of Ccl2, Cxcl9, Cxcl10, Fn1, Ifitm3, and Vcan. The Top 20 markers of this human CCL2-Macrophage subset were mainly expressed by Cxcl9^+^ monocytes, and to a lesser extent by Ifit^+^ TAM C_3. Together, these data indicate that the Cxcl9^+^ monocytes enriched in regressing tumors exist in some human breast tumors and may contribute to the intraepithelial infiltration of CD8 TILs in clinical responders.

## DISCUSSION

During tumor progression, tumor secreted factors drive immune cells to express pro-tumoral phenotypes and support immunosuppression. In successful immunotherapies, this whole system is re-shaped, and myeloid and T cell populations with potent anti-tumoral abilities are generated to efficiently control tumor growth (Bassez et al. 2021; Gubin et al. 2018). Here, we investigated the changes in the CD8^+^ T cell and CD11b^+^ myeloid compartments and their interaction network in the PyMT breast cancer model after treatment using DMXAA, a STING agonist.

Concerning the CD8^+^ T cell compartment, we report CD8^+^ T cells to increasingly express effector memory and stem-cell memory phenotypes after DMXAA treatment. It has been reported that such subpopulations of CD8^+^ T cells at the tumor site are of indicative of effective responses following immunotherapies (van Duikeren et al. 2012; Gattinoni et al. 2011; Principe et al. 2020). Effector memory CD8^+^ T cells produce elevated amounts of cytotoxic factors (GrzB, TNFα, IFNγ) and stem-cell memory cells, most likely originating from a reservoir in draining lymph nodes (Connolly et al. 2021), display prolonged survival.

The generation of such beneficial anti-tumoral CD8^+^ T cells can be directly linked to STING pathway activation in CD8^+^ T cells, as suggested by the type I IFN-related genes we observed in a number of T cell clusters post-DMXAA treatment. Indeed, even though STING signaling strength on T cells must be very precisely controlled not to induce adverse effects (Richter, Paget, and Apetoh 2023), its driving of CD8^+^ T cells into potent anti-tumoral actors is well-established (W. Li et al. 2020; Sen et al. 2019).

Additionally, in parallel, the changes in global signaling pathways that we trigger in the myeloid compartment through activation of the cGAS-STING pathway give rise to differently primed macrophages and monocytes. Type I IFNs and the STING pathway are of particular interest for the anti-tumoral activation of myeloid cells and more especially monocytes (Jneid et al. 2023; Kwart et al. 2022; Lam et al. 2021; Tkach et al. 2022). In our cluster of inflammatory monocytes and interferon-stimulated TAMs, we observed a type I IFNs signature, suggesting their polarization would resemble those found in the mentioned studies. The changes we see in the myeloid compartment suggest loss of pre-existing negative interactions between those cells. It is known that CD8^+^ T cell infiltration in tumor islets and ability to reach their target cells is a crucial element blocking the efficacy of immunotherapies in solid tumors (Mahmoud et al. 2011). Pro-tumoral macrophages can be responsible for such situations as TAMs residing in stromal regions restrain T cells from entering in the islets (Peranzoni et al. 2018). In our setting, it seems that these negative interactions involved TAMs but also DCs have been lifted and replaced by chemotaxis signals.

In such context of immunotherapy-induced myeloid cell activation, the beneficial effects of the therapy become dependent on the presence of these cells. For instance, our team has previously drawn attention to the necessity of reactive TAMs for a proper anti-tumoral response following immunotherapy (Guerin et al. 2019; Thoreau et al. 2015; Weiss et al. 2017).

Still, the exact collaborative mechanisms between activated macrophages and CD8^+^ TILs are not fully understood, as many functions can be carried out by these immune actors in a dynamic manner, a multi-step model comprising various cooperative mechanisms between macrophages and CD8^+^ T cells can be hypothesized (Vermare et al. 2022).

First of all, in this study, our increased cluster of pro-inflammatory monocytes is predicted to produce high levels of ROS and NO in GO analysis, hinting at its participation in the elimination of cancer cells. Indeed, in appropriate conditions, monocytes and macrophages can themselves directly kill tumor cells. This was demonstrated in pre-immunized animals (Bonnotte et al. 2001)and therapeutically-induced tumor control (Nguyen et al. 2018; Thoreau et al. 2015), where these tumoricidal effects relied on NO and ROS secretion as well as TNFα release.

Here, we further highlighted the existence of positive interactions between monocytes and CD8^+^ T cells in the tumors, during the effector phase following DMXAA treatment. Inflammatory monocytes display multiple predicted functions after treatment, with among them chemotaxis towards CD8^+^ T cells. Coherently, we observed better CD8^+^ T cell migration at the tumor site, resulting in better tumor islets infiltration. Interestingly, the subsets of CD8^+^ T cell effector and effector memory, distinguished by CXCR6 expression, were greatly enriched in tumor islets compared to stromal areas, implying they could indeed reach their target tumor cells. Di Pilato et al. have reported the CXCR6^+^ CD8^+^ T cells to remain in perivascular niches in tumors, in close vicinity to CCR7^+^ DCs (Di Pilato et al. 2021). This was established in the context of progressing tumors and additional data insist on the importance to recruit and retain CXCR6^+^ memory effector T cells at the tumor site for proper tumor control upon treatment (Karaki et al. 2021; Matsumura et al. 2008; Muthuswamy et al. 2021), reinforcing our observations that CXCR6^+^CD8^+^ T cells displaying cytotoxic functions migrate into the islets. Our data highlights the CCR2-CCL2 pathway as a potential candidate participating in CD8^+^ T cell attraction to the tumor site.

This pathway is known to be a myeloid-recruiting signaling, as these cells express high levels of the CCR2 receptor and tumors secrete important amounts of CCL2 during its progression, as summarized in (M. Li et al. 2013). However, we here reveal that CCL2 could also be recruiting CD8^+^ T cells post-DMXAA treatment, as the proportion of CD8^+^ T cells expressing CCR2 more than doubles compared to the control condition, and in parallel whole-tumor CCL2 secretion increases during the effector phase. This is intriguing as tumor cells are dying and most of TAMs have disappeared at this time of the response, however inflammatory monocytes signature comprises *Ccl2* as one of the top genes, hinting they could be responsible for this rise of CCL2 levels at the tumor site. Moreover, data from cancer patients suggest that this population is of good prognosis for response to immunotherapy.

Strengthening this chemotaxis hypothesis for monocyte attraction of CD8^+^ T cells in tumor islets are all the other chemotaxis and integrins-mediated interactions enriched in the scRNAseq-derived predictions. Indeed, it is well-established that they are redundancy mechanisms between chemokines, making up for the robustness of these systems. This redundancy makes it difficult to obtain clear results through the blocking of only one chemokine, as we noticed when anti-CCL2 treatment did not prevent DMXAA-induced tumor regression. Indeed, chemokines such as CXCL9, CXCL10 and CXCL12 as well-known to participate in CD8^+^ T cell attraction, as reviewed in (Kohli, Pillarisetty, and Kim 2022; Marcovecchio, Thomas, and Salek-Ardakani 2021). Although we previously showed that TAMs and monocytes in treated mice were capable of producing more CXCL9 after in vitro re-stimulation (Weiss et al. 2017), here our data shows that at the mRNA levels, the expression of CXCL9/10 was decreased post treatment. and the other chemokine of interest, Cxcl12, was not found in our assay. Inversely, our clusters of effector and effector memory CD8^+^ T cells are predicted to further recruit more inflammatory monocytes through the CCR5/CCL5 pathway. This has also been reported after vaccination in a murine lymphoma model where exhausted CD8^+^ T cells attracted CCR5 macrophages to the tumor during the response (van Elsas et al. 2024).

In addition, adhesion molecules such as fibronectin and versican were upregulated in our cluster of monocytes, predicted to bind to CD8^+^ T cells integrins like CD29 (*Itgb1*) and would be involved in the positioning of the T cells in the tumors rather than their recruitment to the inflamed site. Although most probably not directly implicated in the migration of T cells, which is integrin-independent in dense tissues (Fowell and Kim 2021), the binding of those integrins on adhesion molecules could stabilize the cells together for activating long-lasting interactions. Our group has actually previously shown that both CD8^+^ T cells and macrophages heavily infiltrated tumor islets upon DMXAA treatment (Weiss et al. 2017) but a limitation of this study is that we did not manage to see monocyte intra-tumoral localization, which could have shed light on the nature of their interactions with CD8^+^ T cells, depending on their proximity.

In relation to this, and more broadly speaking, it is rather clear that myeloid cell interactions, and functions more generally, depend on the tumor microenvironment area in which they are positioned (Laviron et al. 2022; Nalio Ramos et al. 2022). It would be interesting to determine in which microenvironmental niches these monocytes exert these anti-tumoral functions. We can further hypothesize that the subsets of monocytes and macrophages we detected are localized differently in the tumor architecture, therefore making various cooperation mechanisms non-exclusive, but spatially and dynamically organized inside the tumor. Monocyte cytotoxicity, which can come through NO and ROS secretion, as well as TNFα, could in certain regions, diffuse very locally and eliminate tumor cells, favoring CD8^+^ T cell infiltration in tumor islets.

All in all, this study supports the idea of reprogramming the tumor microenvironment as a whole through therapy-induced signaling changes, to fuel cooperative interaction networks between properly activated immune cells.

## MATERIALS AND METHODS

### Animal studies

#### Animal studies

MMTV-PyMT transgenic mice were maintained at the Cochin Institute SPF animal facility by backcrossing with FVB/NCrl mice (Charles River Laboratories). Transplanted PyMT mice were generated by injecting 1 × 10^6^ cells, freshly isolated from tumor cell suspensions of MMTV-PyMT tumors, into the mammary gland of 8-week-old FVB mice. In some experiments, C57BL/6J mice were transplanted with PyMT-mCherryOVA tumor cell line (M. Krummel, University of California San Francisco, USA). Two weeks after PyMT cells transplantation, mice received a single intraperitoneal (i.p) injection of 5,6-dimethylxanthenone-4-acetic acid (DMXAA) at 23 mg/kg in DMSO (D5817-25MG, Sigma) or 100 µl of PBS 50% DMSO as a control.

Tumor size evolution after DMXAA treatment was calculated relatively to its size at day 0 (d0), when DMXAA was injected. For each mouse, the average size of all measurable tumors was calculated. Animal care complied with all relevant ethical regulations for animal testing and research as per the Federation of European Laboratory Animal Science Associations. All procedures were approved by the French animal experimentation and ethics committee of (Protocol APAFIS no.33813). Sample sizes were chosen to ensure reproducibility of experiments in accordance with the replacement, reduction and refinement principles of animal ethics regulation.

#### Multicolor flow cytometry

Tumor cell suspensions were obtained by mechanical and enzymatic dissociation using DNase I (2.6 x 10^3^ U/ml, 260913-25MU, Roche), liberase (25 μg/ml, 764368, Roche) and hyaluronidase (1 μg/ml, Sigma), for 40 min at 37°C. They were then filtered on 70 µm-filters and rinsed with PBS 2% SVF 1 mM EDTA. Then, red blood cells were lysed using RBC lysis buffer (0-4333-57, Invitrogen) and cell suspensions further filtered on 40 µm-filters. Cell suspensions were rinsed using PBS twice and stained with fluorescent antibodies.

For conventional flow cytometry, cells were first stained with the LIVE/DEAD™ Fixable Blue Dead Cell Stain Kit (#L34962, Invitrogen) at 1/1000 dilution for 10 minutes at 4°C. After this step, cells were rinsed with PBS 2% SVF 1 mM EDTA and stained with 20 µl of antibody cocktail in PBS 2% SVF 1 mM EDTA with 5 µg/ml anti-CD16/32 (clone 93, *#101302,* Biolegend) (see Table XXX for references and dilution of antibodies) for 20 minutes at 4°C with gentle shaking. After these steps, cells were washed with PBS 2% SVF 1 mM EDTA and fixed with 100µl PFA 4% for 10 minutes at room temperature. Cells were washed and resuspended in PBS 2% SVF 1mM EDTA for flow cytometer acquisition on a BD LSR Fortessa flow cytometer (BD Biosciences, USA).

For spectral flow cytometry, tumor cell suspensions were stained with LIVE/DEAD™ Fixable Aqua Stain (*L34966, Invitrogen*) during 30 min, at 4°C, with gentle shaking . Cells were then washed in PBS 2 mM EDTA 0.2 % BSA. Fc receptors were blocked with 5 µg/ml anti-CD16/32 (clone 2.4G2, *#BE0307,* BioXCell) for 10 min, at 4°C. For surface staining, cells were incubated with antibodies in PBS containing 5 µg/ml anti-CD16/32, for 30 min, at 4°C with gentle shaking. After one wash in PBS 2 mM EDTA 0.2 % BSA, cells were fixed in 2% PFA for 10 min at 4°C. Cells were then permeabilized using the Foxp3/Transcription Factor Staining Buffer Set (# 00-5523-00, eBioscience), according to manufacturer’s procedures. Antibody staining was performed overnight at 4°C in permeabilization solution, before extensive washing with PBS 2 mM EDTA 0.2 % BSA.

T cell subsets were analyzed using the appropriate combination of the following antibodies (Table 1): CD45-BUV395, CD8-BV615, PD1-BV650, CD44-BV786, CD62L-APC, TIM-3-PE, TCR-β-AF700, TCF-1-BV421, Foxp3-FITC, Tbet-PEVio615 and GranzymeB-APC/Fire750. Myeloid cell subsets were analyzed using CD45-BUV395, Ly6G-BV711, CD11b-PE-Cy7, Ly6C-PerCP-Cy5.5. Data were acquired the following day on an Aurora spectral flow cytometer (Cytek, USA).

All flow cytometry data were analyzed using FlowJo software 10.9.0 (*BD Biosciences*).

#### Immunofluorescence on tumor slices

Fresh tumor samples were fixed overnight at 4 °C using 0.075 M lysine, 0.37 M sodium phosphate 2% paraformaldehyde, periodate–lysine–paraformaldehyde (PLP) and preserved in PBS 0.01% Na-Azide at 4°C. Tumor slices were prepared as described in [Guerin2019] and stained for immunofluorescence. Non-specific binding sites on the tissue slices were blocked with PBS 0.5% BSA 0.05% TRITON-X for 10 minutes. The slices were then stained with conjugated antibodies for 15 minutes at 37°C. The following anti-mouse antibodies were used (Table X). After staining, tissue slices were rinsed with PBS 0,01% Tween and mounted with Antifade Vectashield (Vector Labs) and imaged using an inverted spinning disk confocal microscope (Olympus IXplore, CSU-W1 T1; Yokogawa) with a 20x objective using the cellSens acquiring software (IX3-DSU, IX83F).

#### Live Imaging

Tumors from DMXAA or control mice were collected at day4 to 6 after treatment and were used to prepare 400 µm-thick slices. Slices were kept in PBS supplemented with 2% SVF at 4°C until use. Slices were put 30 minutes in an incubator at 37°C with 5% CO_2_ before staining and imaging. In the experiments with CCR2 inhibition, the antagonist (ref. RS504393 from MedChem Express) was added in 2 ml at the final concentration of 1 µM during this incubation time and the slices were rinsed before the staining step. For the staining, slices were incubated with fluorochrome-conjugated antibodies for 15 minutes at 37°C, then, rinsed and gently attached to the bottom of a 50 mm-Petri dish using a surgical glue covered in PBS 1% SVF 1 mM Na-pyruvate. The dishes containing the stained slice were imaged on an immersion spinning disk confocal microscope (DM6000 FS, Leica with a CSU-X1 Yokogawa head) equipped with a thermostatic chamber kept at 37°C with 5% CO_2_. During time lapse acquisition, the slices were continuously perfused with PBS 1% SVF 1 mM Na-pyruvate at very low speed. Images were taken every 45 seconds for 30 minutes for CD8 and CXCR6; EpCAM and gp38 images were only taken every 10 time points to avoid bleaching. Z-stack images of 70µm were taken to track CD8^+^ cells in the 3D microenvironment. Time lapse videos were analyzed using ImageJ software and the TrackMate plugin. CD8^+^ T cell average speed and traveled distances were represented.

#### Transcriptomic analysis

Tumor cell suspensions were stained with xx Abs before sorting with a FACS ARIA III (BD Biosciences) using a 85µm nozzle at 45psi. Cells were sorted into chilled eppendorf tubes containing 15 µL of PBS-50% FBS and immediately used for subsequent processing. Single-cell Gel Bead-In-EMulsions (GEMs) were generated using a Chromium Controller instrument (10x Genomics). Sequencing libraries were prepared using Chromium Single Cell 3’ Reagent Kits v3.1 (10x Genomics), according to the manufacturer’s instructions. Briefly, GEM-RT was performed in a thermal cycler: 53°C for 45 min, 85°C for 5 min. Post GEM-RT Cleanup using DynaBeads MyOne Silane Beads was followed by cDNA amplification (98°C for 3 min, cycled 11 x 98°C for 15 s, 67°C for 20s, 72°C for1 min, and 72°C 1 min). After a cleanup with SPRIselect Reagent Kit and fragment size estimation with High Sensitivity HS DNA kit runned on 2100 Bioanalyzer (Agilent), the libraries were constructed by performing the following steps: fragmentation, end-repair, A-tailing, SPRIselect cleanup, adaptor ligation, SPRIselect cleanup, sample index PCR, and SPRIselect size selection.

The fragment size estimation of the resulting libraries was assessed with High Sensitivity HS DNA kit runned on 2100 Bioanalyzer (Agilent) and quantified using the Qubit™ dsDNA High Sensitivity HS assay (ThermoFisher Scientific). Libraries were then sequenced by pair with a High Output flowcel using an Illumina Nextseq 500 with the following mode: 28 bp (10X Index + UMI), 8 bp (i7 Index) and 91 bp (Read 2).

The sequencing data was processed into transcript count tables with the Cell Ranger Single Cell Software Suite 1.3.1 by 10X Genomics (http://10xgenomics.com/). Raw base call files from the Nextseq 500 were demultiplexed with the *cellranger mkfastq* pipeline into library-specific FASTQ files. The FASTQ files for each library were then processed independently with the *cellranger count* pipeline. This pipeline used STAR to align cDNA reads to the Mus musculus transcriptome (Sequence: GRCm38, Annotation: Gencode v25). Once aligned, barcodes associated with these reads – cell identifiers and Unique Molecular Identifiers (UMIs), underwent filtering and correction. Reads associated with retained barcodes were quantified and used to build a transcript count table. Resulting data for each sample were then aggregated using the *cellranger aggr* pipeline, which performed a between-sample normalization step and concatenated the two transcript count tables. Post-aggregation, the mapped data was processed and analyzed as described below.

After sequencing, a primary analysis based on the 10x Genomics Cell Ranger 4.0.0 software (Zheng et al. 2017) was applied to demultiplex, control the quality of the raw data and assign each read to one gene and one cell. Obtained filtered_feature_bc_matrix repertory was then analysed with R version 3.6.3 and the Seurat_3.2.2 (Stuart et al. 2019) package. Figures were prepared using the ggplot2_3.3.2 package (Gómez Rubio 2017). Pre-filters: each gene has to be shared by at least 3 cells to be conserved, each cell must express between 200 and 3000 genes with less than 5% of mitochondrion genes. Integration: The integration of two datasets was performed with the FindIntegrationAnchors and IntegrateData functions. Once the dataset ready (standard pipeline), the Find Markers function was applied to get specific markers by cluster and specific marker for each sample, for a given cluster. All functions were mostly used with default parameters, and R script are available.

Cell-cell interactions and communication were studied using CellPhoneDB v3.0.0 [PMID: 32103204], NATMI [PMID: 33024107] and NicheNet v1.0.0 [PMID: 31819264] with default parameters.

Comparisons with the GSE169246 GEO dataset from Zhang et al. [PMID: 34653365] were performed by label transfer using FindTransferAnchors and TransferData from Seurat library (v5.0.1) in R (v4.3.2).

#### Cytokine measurements in tumor samples

Tumor slices were cultivated on a Millicell® Standing Cell Culture Inserts filter (#PICM01250, Millipore) for 48 h at 37°C with 5% CO2 in DMEM 10 % FBS, 10 x 10^3^ UI/ml Penicilin, 10 x 10^3^ µg/ml Streptomycin, 100 mM Na Pyruvate.

The concentrations of 13 mouse cytokines (interferons α, β and γ, IL-1β, IL-6, IL-10, IL-12, TNF-α and GM-CSF) and chemokines (CCL2/MCP-1, CCL5/RANTES, CXCL1/KC, CXCL10/IP-10) were quantified using the LEGENDplex™ Mouse Anti-Virus Response Cytokine Panel Standard (#740621, BioLegend), according to the manufacturer’s protocol. All samples were measured in duplicates. Briefly, supernatants and standards were diluted in the LEGENDplex™ assay buffer and then incubated in V-bottom 96 well plates with capture antibody-conjugated beads for 2 h, at RT, with a shaking at 800 rpm. Beads were washed 5 min at 1050 rpm, RT, samples were incubated with biotinylated detection antibodies for 60 min, and finally with streptavidin-PE for 30 min. Acquisition was performed on BD LSR Fortessa flow cytometer (*BD Biosciences, USA*) using BD™ High Throughput Sampler (*HTS, BD Biosciences, USA*).

## REFERENCES

1. Barras, David, Eleonora Ghisoni, Johanna Chiffelle, Angela Orcurto, Julien Dagher, Noémie Fahr, Fabrizio Benedetti, et al. 2024. ‘Response to Tumor-Infiltrating Lymphocyte Adoptive Therapy Is Associated with Preexisting CD8+ T-Myeloid Cell Networks in Melanoma’. Science Immunology 9(92): eadg7995. doi:10.1126/sciimmunol.adg7995.

2. Bassez, Ayse, Hanne Vos, Laurien Van Dyck, Giuseppe Floris, Ingrid Arijs, Christine Desmedt, Bram Boeckx, et al. 2021. ‘A Single-Cell Map of Intratumoral Changes during Anti-PD1 Treatment of Patients with Breast Cancer’. Nature Medicine 27(5): 820–32. doi:10.1038/s41591-021-01323-8.

3. Bonnotte, Bernard, Nicolas Larmonier, Nathalie Favre, Annie Fromentin, Monique Moutet, Monique Martin, Sandeep Gurbuxani, et al. 2001. ‘Identification of Tumor-Infiltrating Macrophages as the Killers of Tumor Cells After Immunization in a Rat Model System1’. The Journal of Immunology 167(9): 5077–83. doi:10.4049/jimmunol.167.9.5077.

4. Connolly, Kelli A., Manik Kuchroo, Aarthi Venkat, Achia Khatun, Jiawei Wang, Ivana William, Noah I. Hornick, et al. 2021. ‘A Reservoir of Stem-like CD8+ T Cells in the Tumor-Draining Lymph Node Preserves the Ongoing Antitumor Immune Response’. Science Immunology 6(64): eabg7836. doi:10.1126/sciimmunol.abg7836.

5. Dangaj, Denarda, Marine Bruand, Alizée J. Grimm, Catherine Ronet, David Barras, Priyanka A. Duttagupta, Evripidis Lanitis, et al. 2019. ‘Cooperation between Constitutive and Inducible Chemokines Enables T Cell Engraftment and Immune Attack in Solid Tumors’. Cancer Cell 35(6): 885–900.e10. doi:10.1016/j.ccell.2019.05.004.

6. Di Pilato, Mauro, Raphael Kfuri-Rubens, Jasper N. Pruessmann, Aleksandra J. Ozga, Marius Messemaker, Bruno L. Cadilha, Ramya Sivakumar, et al. 2021. ‘CXCR6 Positions Cytotoxic T Cells to Receive Critical Survival Signals in the Tumor Microenvironment’. Cell 184(17): 4512–4530.e22. doi:10.1016/j.cell.2021.07.015.

7. van Duikeren, Suzanne, Marieke F. Fransen, Anke Redeker, Brigitte Wieles, Gerard Platenburg, Willem-Jan Krebber, Ferry Ossendorp, Cornelis J. M. Melief, and Ramon Arens. 2012. ‘Vaccine-Induced Effector-Memory CD8+ T Cell Responses Predict Therapeutic Efficacy against Tumors’. The Journal of Immunology 189(7): 3397–3403. doi:10.4049/jimmunol.1201540.

8. van Elsas, Marit J., Jim Middelburg, Camilla Labrie, Jessica Roelands, Gaby Schaap, Marjolein Sluijter, Ruxandra Tonea, et al. 2024. ‘Immunotherapy-Activated T Cells Recruit and Skew Late-Stage Activated M1-like Macrophages That Are Critical for Therapeutic Efficacy’. Cancer Cell 42(6): 1032–1050.e10. doi:10.1016/j.ccell.2024.04.011.

9. Fowell, Deborah J., and Minsoo Kim. 2021. ‘The Spatio-Temporal Control of Effector T Cell Migration’. Nature Reviews. Immunology 21(9): 582–96. doi:10.1038/s41577-021-00507-0.

10. Gattinoni, Luca, Enrico Lugli, Yun Ji, Zoltan Pos, Chrystal M. Paulos, Máire F. Quigley, Jorge R. Almeida, et al. 2011. ‘A Human Memory T Cell Subset with Stem Cell-like Properties’. Nature Medicine 17(10): 1290–97. doi:10.1038/nm.2446.

11. Gómez Rubio, Virgilio. 2017. ‘Book Review: Ggplot2 – Elegant Graphics for Data Analysis (2nd Edition)’. Journal of statistical software 77. doi:10.18637/jss.v077.b02.

12. Gubin, Matthew M., Ekaterina Esaulova, Jeffrey P. Ward, Olga N. Malkova, Daniele Runci, Pamela Wong, Takuro Noguchi, et al. 2018. ‘High-Dimensional Analysis Delineates Myeloid and Lymphoid Compartment Remodeling during Successful Immune-Checkpoint Cancer Therapy’. Cell 175(4): 1014–1030.e19. doi:10.1016/j.cell.2018.09.030.

13. Guerin, Marion V., Fabienne Regnier, Vincent Feuillet, Lene Vimeux, Julia M. Weiss, Georges Bismuth, Gregoire Altan-Bonnet, et al. 2019. ‘TGFβ Blocks IFNα/β Release and Tumor Rejection in Spontaneous Mammary Tumors’. Nature Communications 10(1): 4131. doi:10.1038/s41467-019-11998-w.

14. Jneid, Bakhos, Aurore Bochnakian, Caroline Hoffmann, Fabien Delisle, Emeline Djacoto, Philémon Sirven, Jordan Denizeau, et al. 2023. ‘Selective STING Stimulation in Dendritic Cells Primes Antitumor T Cell Responses’. Science Immunology 8(79): eabn6612. doi:10.1126/sciimmunol.abn6612.

15. Karaki, Soumaya, Charlotte Blanc, Thi Tran, Isabelle Galy-Fauroux, Alice Mougel, Estelle Dransart, Marie Anson, et al. 2021. ‘CXCR6 Deficiency Impairs Cancer Vaccine Efficacy and CD8+ Resident Memory T-Cell Recruitment in Head and Neck and Lung Tumors’. Journal for Immunotherapy of Cancer 9(3): e001948. doi:10.1136/jitc-2020-001948.

16. Kohli, Karan, Venu G. Pillarisetty, and Teresa S. Kim. 2022. ‘Key Chemokines Direct Migration of Immune Cells in Solid Tumors’. Cancer Gene Therapy 29(1): 10–21. doi:10.1038/s41417-021-00303-x.

17. Kwart, Dylan, Jing He, Subhashini Srivatsan, Clarissa Lett, Jacquelynn Golubov, Erin M. Oswald, Patrick Poon, et al. 2022. ‘Cancer Cell-Derived Type I Interferons Instruct Tumor Monocyte Polarization’. Cell Reports 41(10): 111769. doi:10.1016/j.celrep.2022.111769.

18. Lam, Khiem C., Romina E. Araya, April Huang, Quanyi Chen, Martina Di Modica, Richard R. Rodrigues, Amélie Lopès, et al. 2021. ‘Microbiota Triggers STING-Type I IFN-Dependent Monocyte Reprogramming of the Tumor Microenvironment’. Cell 184(21): 5338–5356.e21. doi:10.1016/j.cell.2021.09.019.

19. Laviron, Marie, Maxime Petit, Eléonore Weber-Delacroix, Alexis J. Combes, Arjun Rao Arkal, Sandrine Barthélémy, Tristan Courau, et al. 2022. ‘Tumor-Associated Macrophage Heterogeneity Is Driven by Tissue Territories in Breast Cancer’. Cell Reports 39(8): 110865. doi:10.1016/j.celrep.2022.110865.

20. Li, Ming, Deborah A. Knight, Linda A Snyder, Mark J. Smyth, and Trina J. Stewart. 2013. ‘A Role for CCL2 in Both Tumor Progression and Immunosurveillance’. Oncoimmunology 2(7): e25474. doi:10.4161/onci.25474.

21. Li, Wenwen, Lu Lu, Juanjuan Lu, Xinran Wang, Chao Yang, Jingsi Jin, Lingling Wu, et al. 2020. ‘cGAS-STING-Mediated DNA Sensing Maintains CD8+ T Cell Stemness and Promotes Antitumor T Cell Therapy’. Science Translational Medicine 12(549): eaay9013. doi:10.1126/scitranslmed.aay9013.

22. Mahmoud, Sahar M. A., Emma Claire Paish, Desmond G. Powe, R. Douglas Macmillan, Matthew J. Grainge, Andrew H. S. Lee, Ian O. Ellis, and Andrew R. Green. 2011. ‘Tumor-Infiltrating CD8+ Lymphocytes Predict Clinical Outcome in Breast Cancer’. Journal of Clinical Oncology: Official Journal of the American Society of Clinical Oncology 29(15): 1949–55. doi:10.1200/JCO.2010.30.5037.

23. Mantovani, Alberto, Federica Marchesi, Alberto Malesci, Luigi Laghi, and Paola Allavena. 2017. ‘Tumour-Associated Macrophages as Treatment Targets in Oncology’. Nature Reviews Clinical Oncology 14(7): 399–416. doi:10.1038/nrclinonc.2016.217.

24. Mantovani, Alberto, Silvano Sozzani, Massimo Locati, Paola Allavena, and Antonio Sica. 2002. ‘Macrophage Polarization: Tumor-Associated Macrophages as a Paradigm for Polarized M2 Mononuclear Phagocytes’. Trends in Immunology 23(11): 549–55. doi:10.1016/S1471-4906(02)02302-5.

25. Marcovecchio, Paola Marie, Graham Thomas, and Shahram Salek-Ardakani. 2021. ‘CXCL9-Expressing Tumor-Associated Macrophages: New Players in the Fight against Cancer’. Journal for Immunotherapy of Cancer 9(2): e002045. doi:10.1136/jitc-2020-002045.

26. Matsumura, Satoko, Baomei Wang, Noriko Kawashima, Steve Braunstein, Michelle Badura, Thomas O. Cameron, James S. Babb, et al. 2008. ‘Radiation-Induced CXCL16 Release by Breast Cancer Cells Attracts Effector T Cells’. Journal of Immunology (Baltimore, Md.: 1950) 181(5): 3099–3107. doi:10.4049/jimmunol.181.5.3099.

27. Mulder, Kevin, Amit Ashok Patel, Wan Ting Kong, Cécile Piot, Evelyn Halitzki, Garett Dunsmore, Shabnam Khalilnezhad, et al. 2021. ‘Cross-Tissue Single-Cell Landscape of Human Monocytes and Macrophages in Health and Disease’. Immunity 54(8): 1883–1900.e5. doi:10.1016/j.immuni.2021.07.007.

28. Muthuswamy, Ravikumar, Aj Robert McGray, Sebastiano Battaglia, Wenjun He, Anthony Miliotto, Cheryl Eppolito, Junko Matsuzaki, et al. 2021. ‘CXCR6 by Increasing Retention of Memory CD8+ T Cells in the Ovarian Tumor Microenvironment Promotes Immunosurveillance and Control of Ovarian Cancer’. Journal for Immunotherapy of Cancer 9(10): e003329. doi:10.1136/jitc-2021-003329.

29. Nalio Ramos, Rodrigo, Yoann Missolo-Koussou, Yohan Gerber-Ferder, Christian P. Bromley, Mattia Bugatti, Nicolas Gonzalo Núñez, Jimena Tosello Boari, et al. 2022. ‘Tissue-Resident FOLR2+ Macrophages Associate with CD8+ T Cell Infiltration in Human Breast Cancer’. Cell 185(7): 1189–1207.e25. doi:10.1016/j.cell.2022.02.021.

30. Nguyen, Andrew, Louisa Ho, Samuel T. Workenhe, Lan Chen, Jonathan Samson, Scott R. Walsh, Jonathan Pol, Jonathan L. Bramson, and Yonghong Wan. 2018. ‘HDACi Delivery Reprograms Tumor-Infiltrating Myeloid Cells to Eliminate Antigen-Loss Variants’. Cell Reports 24(3): 642–54. doi:10.1016/j.celrep.2018.06.040.

31. Peranzoni, Elisa, Jean Lemoine, Lene Vimeux, Vincent Feuillet, Sarah Barrin, Chahrazade Kantari-Mimoun, Nadège Bercovici, et al. 2018. ‘Macrophages Impede CD8 T Cells from Reaching Tumor Cells and Limit the Efficacy of Anti-PD-1 Treatment’. Proceedings of the National Academy of Sciences of the United States of America 115(17): E4041–50. doi:10.1073/pnas.1720948115.

32. Principe, Nicola, Joel Kidman, Siting Goh, Caitlin M. Tilsed, Scott A. Fisher, Vanessa S. Fear, Catherine A. Forbes, et al. 2020. ‘Tumor Infiltrating Effector Memory Antigen-Specific CD8+ T Cells Predict Response to Immune Checkpoint Therapy’. Frontiers in Immunology 11: 584423. doi:10.3389/fimmu.2020.584423.

33. Richter, Fabian, Christophe Paget, and Lionel Apetoh. 2023. ‘STING-Driven Activation of T Cells: Relevance for the Adoptive Cell Therapy of Cancer’. Cell Stress 7(11): 95–104. doi:10.15698/cst2023.11.291.

34. Sen, Triparna, B. Leticia Rodriguez, Limo Chen, Carminia M. Della Corte, Naoto Morikawa, Junya Fujimoto, Sandra Cristea, et al. 2019. ‘Targeting DNA Damage Response Promotes Antitumor Immunity through STING-Mediated T-Cell Activation in Small Cell Lung Cancer’. Cancer Discovery 9(5): 646–61. doi:10.1158/2159-8290.CD-18-1020.

35. Stuart, Tim, Andrew Butler, Paul Hoffman, Christoph Hafemeister, Efthymia Papalexi, William M. Mauck, Yuhan Hao, et al. 2019. ‘Comprehensive Integration of Single-Cell Data’. Cell 177(7): 1888–1902.e21. doi:10.1016/j.cell.2019.05.031.

36. Thoreau, Maxime, HweiXian Leong Penny, KarWai Tan, Fabienne Regnier, Julia Miriam Weiss, Bernett Lee, Ludger Johannes, et al. 2015. ‘Vaccine-Induced Tumor Regression Requires a Dynamic Cooperation between T Cells and Myeloid Cells at the Tumor Site’. Oncotarget 6(29): 27832–46. doi:10.18632/oncotarget.4940.

37. Tkach, Mercedes, Jessie Thalmensi, Eleonora Timperi, Paul Gueguen, Nathalie Névo, Eleonora Grisard, Philemon Sirven, et al. 2022. ‘Extracellular Vesicles from Triple Negative Breast Cancer Promote Pro-Inflammatory Macrophages Associated with Better Clinical Outcome’. Proceedings of the National Academy of Sciences of the United States of America 119(17): e2107394119. doi:10.1073/pnas.2107394119.

38. Vermare, Anaïs, Marion V. Guérin, Elisa Peranzoni, and Nadège Bercovici. 2022. ‘Dynamic CD8+ T Cell Cooperation with Macrophages and Monocytes for Successful Cancer Immunotherapy’. Cancers 14(14): 3546. doi:10.3390/cancers14143546.

39. Weiss, Julia M., Marion V. Guérin, Fabienne Regnier, Gilles Renault, Isabelle Galy-Fauroux, Lene Vimeux, Vincent Feuillet, et al. 2017. ‘The STING Agonist DMXAA Triggers a Cooperation between T Lymphocytes and Myeloid Cells That Leads to Tumor Regression’. Oncoimmunology 6(10): e1346765. doi:10.1080/2162402X.2017.1346765.

40. Zhang, Yuanyuan, Hongyan Chen, Hongnan Mo, Xueda Hu, Ranran Gao, Yahui Zhao, Baolin Liu, et al. 2021. ‘Single-Cell Analyses Reveal Key Immune Cell Subsets Associated with Response to PD-L1 Blockade in Triple-Negative Breast Cancer’. Cancer Cell 39(12): 1578–1593.e8. doi:10.1016/j.ccell.2021.09.010.

41. Zheng, Grace X. Y., Jessica M. Terry, Phillip Belgrader, Paul Ryvkin, Zachary W. Bent, Ryan Wilson, Solongo B. Ziraldo, et al. 2017. ‘Massively Parallel Digital Transcriptional Profiling of Single Cells’. Nature Communications 8: 14049. doi:10.1038/ncomms14049.

