## Supplemental Figures for "Monocyte-derived macrophages promote intraepithelial infiltration of effector memory CD8^+^ T cells in tumors regressing after STING agonist treatment"

Fig. S1

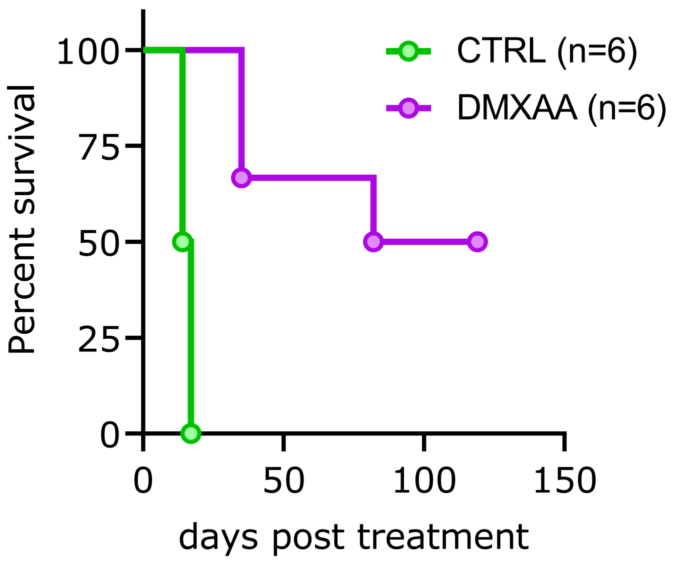

CTRL (n=6)
DMXAA (n=6)
Percent survival
100
75
50
25
0
0
50
100
150
days post treatment

**Figure Supp 1 – Prolonged survival of mice when treated with DMXAA**

Representative experiment of FvB mice transplanted with PyMT tumors and treated with one i.p injection of DMXAA (purple graph) or PBS 50% DMSO as a control (green graph) and their survival (controls n= 6, DMXAA = 6, from 1 experiment).

Fig. S2

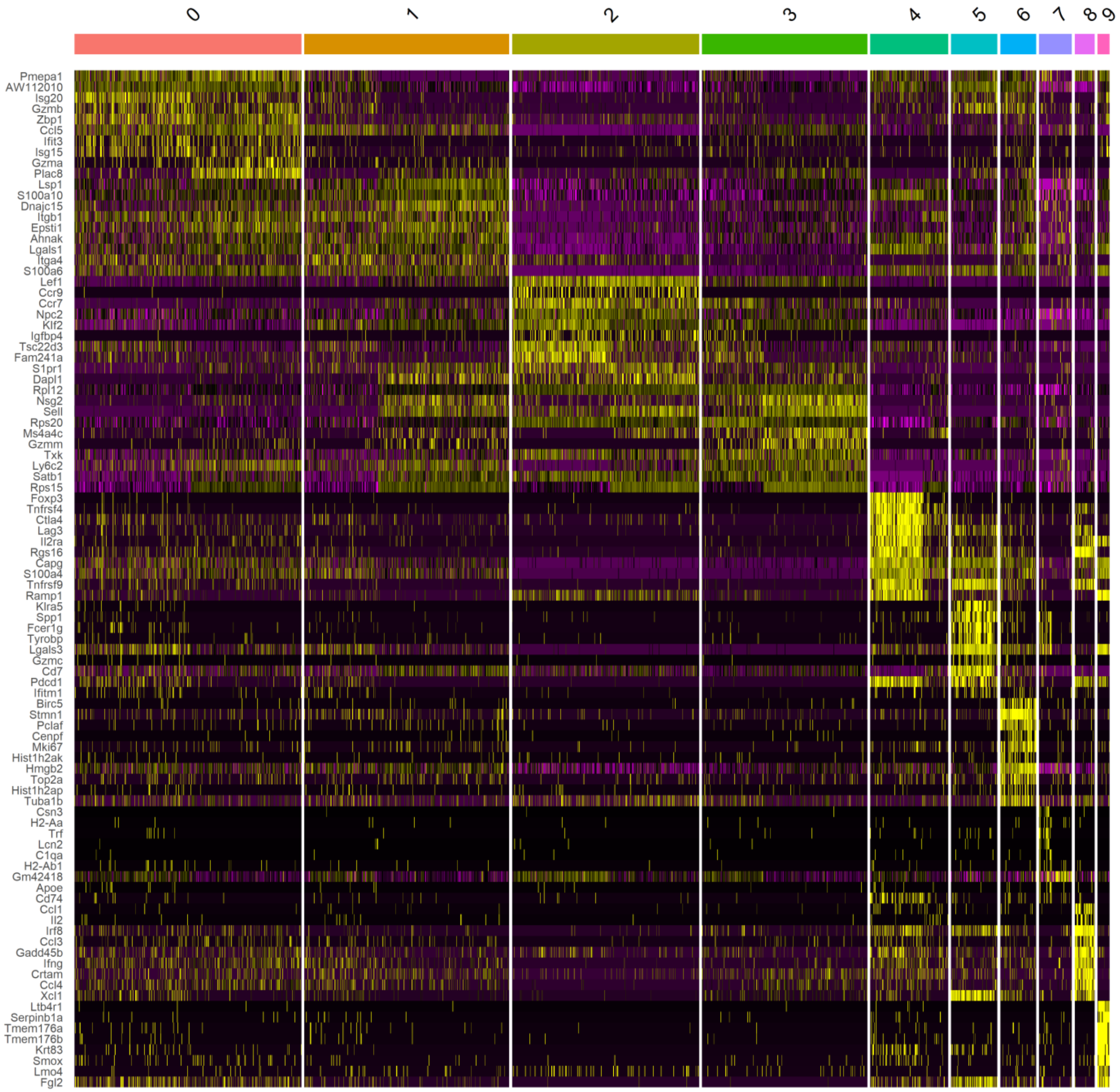
0
1
2
3
4
5
6
7
8
9
Pmepa1
AW112010
Isg20
Gzmb
Zbp1
Ccl5
Ifit3
Isg15
Gzma
Plac8
Lsp1
S100a10
Dnajc15
Itgb1
Epsti1
Ahnak
Lgals1
Itga4
S100a6
Lef1
Ccr9
Ccr7
Npc2
Klf2
Igfbp4
Tsc22d3
Fam241a
S1pr1
Dapl1
Rpl12
Nsg2
Sell
Rps20
Ms4a4c
Gzmm
Txk
Ly6c2
Satb1
Rps15
Foxp3
Tnfrsf4
Ctla4
Lag3
Il2ra
Rgs16
Capg
S100a4
Tnfrsf9
Ramp1
Klra5
Spp1
Fcer1g
Tyrobp
Lgals3
Gzmc
Cd7
Pdcd1
Ifitm1
Birc5
Stmn1
Pclaf
Cenpf
Mki67
Hist1h2ak
Hmgb2
Top2a
Hist1h2ap
Tuba1b
Csn3
H2-Aa
Trf
Lcn2
C1qa
H2-Ab1
Gm42418
Apoe
Cd74
Ccl1
Il2
Irf8
Ccl3
Gadd45b
Ifng
Crtam
Ccl4
Xcl1
Ltb4r1
Serpinb1a
Tmem176a
Tmem176b
Krt83
Smox
Lmo4
Fgl2

**Figure Supp 2 – Top 15 genes for CD8$^{+}$ myeloid clusters**

Heatmap with the top 15 genes for each cluster of CD8$^{+}$ T cells clusters, after scRNAseq analysis. DMXAA, n=26 and controls n=6, merged by group for one experiment.

Fig. S3

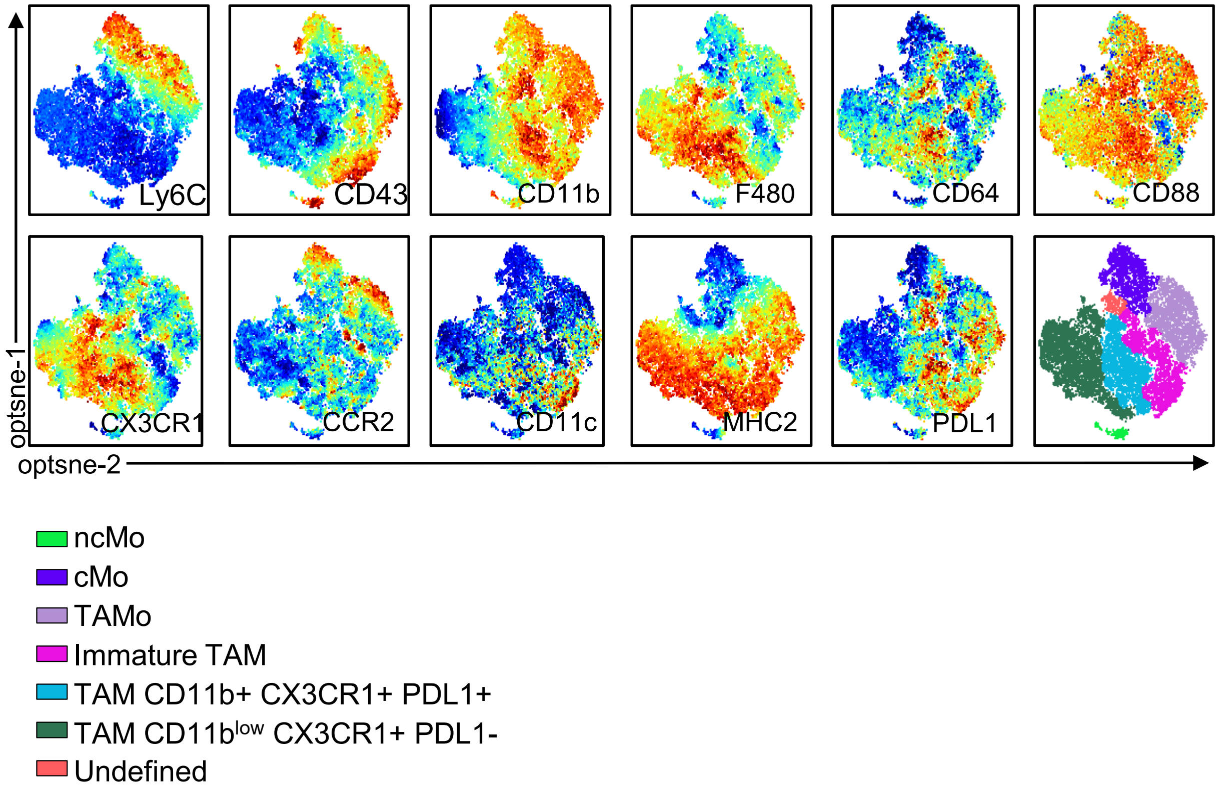
Ly6C
CD43
CD11b
F480
CD64
CD88
CX3CR1
CCR2
CD11c
MHC2
PDL1
optsne-1
optsne-2
ncMo
cMo
TAMo
Immature TAM
TAM CD11b+ CX3CR1+ PDL1+
TAM CD11b$^{low}$ CX3CR1+ PDL1-
Undefined

**Figure Supp 3 – Expression patterns of spectral markers by myeloid population**

Monocyte and macrophages diversity was assessed by spectral flow cytometry, based on their expression of several surface markers: Ly6C, CD43, CD11b, F4/80, CD64, CD88, $CX_3CR1$, CCR2, CD11c, MHC-II and PDL1. The tSNE represents for each marker, the intensity expressed by the analyzed cells, in a red to blue gradient, where red in the high levels and blue the low levels of expression.

### Fig. S4

Exp. 1

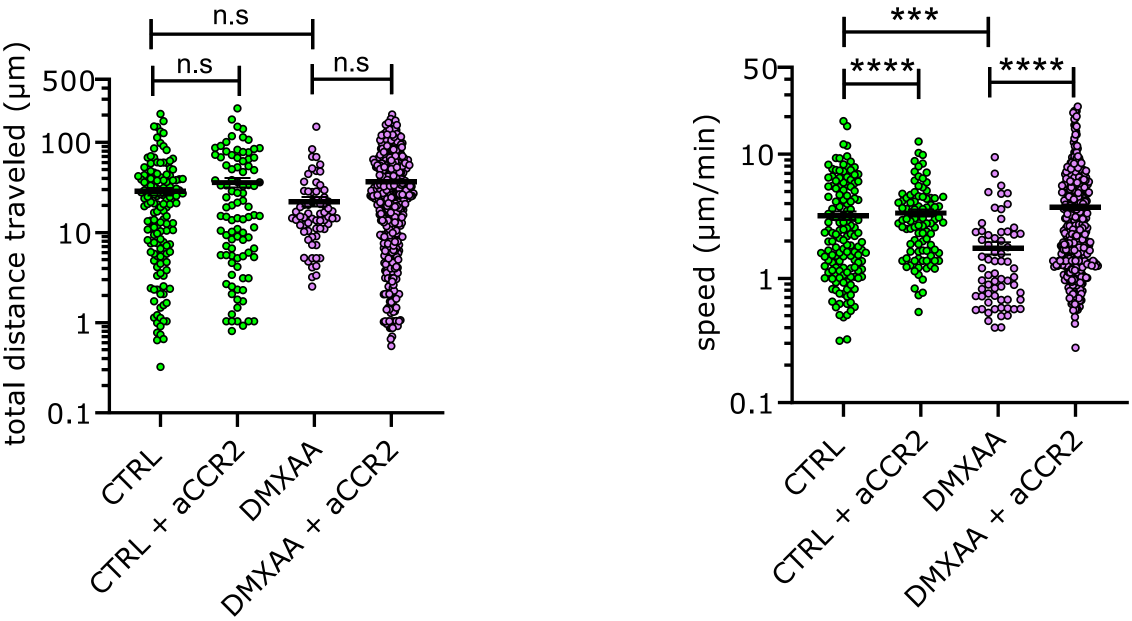


Exp. 2

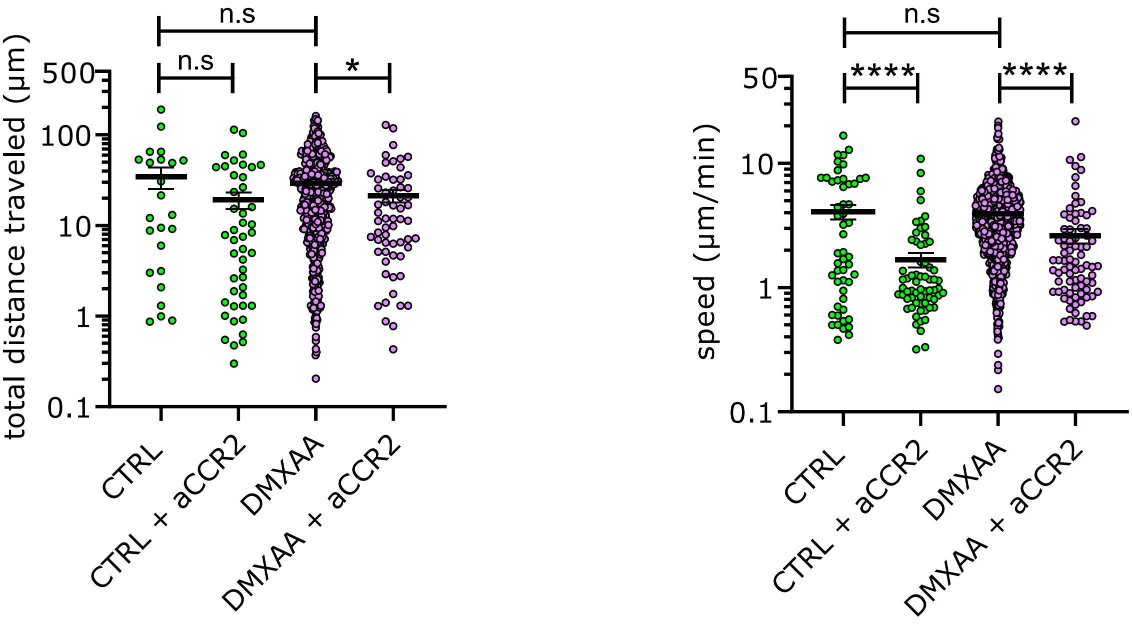

**Figure Supp 4 – *in vitro* CD8$^+$ T cell migration in thick tumor slices when blocking CCR2**

Live imaging movies following CD8$^+$ cells migration in slices of tumors with a pre-incubation of slices with a CCR2 antagonist were made. Here, the quantification of CD8$^+$ cell mean speed (µm/min) and distance traveled (µm) quantifications, from control or treated mice (day 4 post-DMXAA). DMXAA, n=1 tumor and controls n=1 tumor, for each experiment.
Multiple comparison non-parametric Kruskal-Wallis statistical test with Dunn correction, for each experiment separately. **** $p < 0.0001$.

### Fig. S5

A

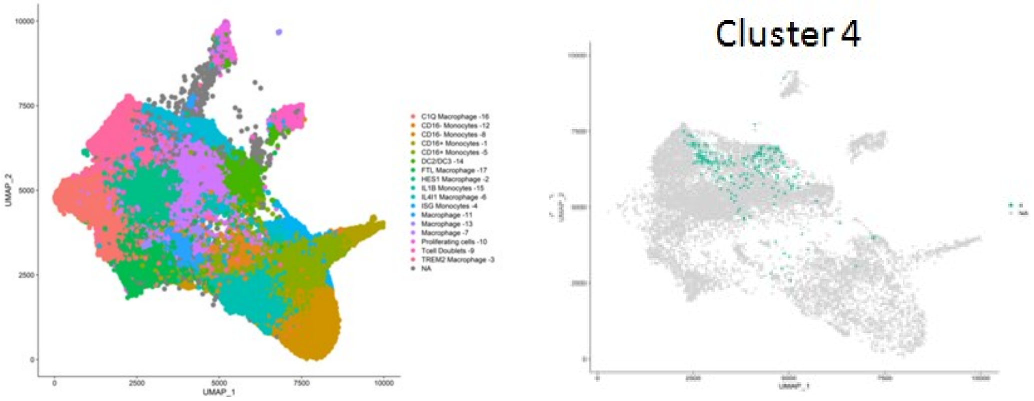
UMAP_1
UMAP_2
0
2500
5000
7500
10000
C1Q Macrophage -16
CD16- Monocytes -12
CD16- Monocytes -8
CD16+ Monocytes -1
CD16+ Monocytes -5
DC2/DC3 -14
FTL Macrophage -17
HES1 Macrophage -2
IL1B Monocytes -15
IL4I1 Macrophage -6
ISG Monocytes -4
Macrophage -11
Macrophage -13
Macrophage -7
Proliferating cells -10
Tcell Doublets -9
TREM2 Macrophage -3
NA
Cluster 4

B

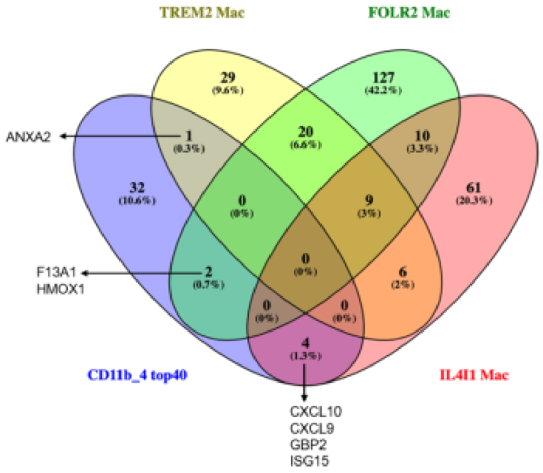
TREM2 Mac
FOLR2 Mac
CD11b_4 top40
IL4I1 Mac
29
(9.6%)
127
(42.2%)
1
(0.3%)
20
(6.6%)
10
(3.3%)
32
(10.6%)
0
(0%)
9
(3%)
61
(20.3%)
0
(0%)
2
(0.7%)
6
(2%)
0
(0%)
0
(0%)
4
(1.3%)
ANXA2
F13A1
HMOX1
CXCL10
CXCL9
GBP2
ISG15

**Figure Supp 5 – Overlap of our cluster 4 of inflammatory monocytes onto the MoMac VERSE, by Mulder et al. 2021.**

**(A)**The MoMac-VERSE with the projection of the clusters, from Mulder et al. 2021 (on the left). Our cluster 4 is projected onto the MoMac-VERSE and is split between TREM2 Mac, FOLR2 Mac and IL4i1 Mac (on the right). **(B)** Venn diagram representing the number of genes from the clusters' top genes, crossing between populations. CXCL9, CXCL10 are in common between our cluster 4 and the IL4i1 macrophages from the MoMac-VERSE.
